# Towards a general Detector of terrestrial Arthropods in Natural backgrounds

**DOI:** 10.64898/2026.05.06.723207

**Authors:** Edgar Remy, Axel Carlier, Elodie Massol, Rahim Kacimi, Alexis S. Chaine, Maxime Cauchoix

## Abstract

Widespread arthropod declines pose risks to ecosystem functioning and agriculture. Assessing this decline or potential remediation implies the need for standardized and scalable population monitoring. Image-based methods, including camera traps and citizen science programs, are increasingly used, but the volume of data collected requires automated analysis. Robust arthropod detection is essential for individual counting or fine-grained classification, yet current datasets and algorithms do not address the vast morphological diversity across arthropod species and often overlook the variety of photographic contexts, such as differences in background, lighting, and image composition, in which arthropods are captured.

To address this gap, we developed an arthropod detection dataset, covering all terrestrial families present in France with available validated images on the iNaturalist platform (749 families). To achieve this, we employed an iterative workflow in which a YOLOv11 model pre-annotated images — using one representative species per family— followed by manual correction and model retraining. Repeating this process progressively reduced annotation effort and improved model accuracy.

The final outcome consists of a publicly available curated detection dataset and a robust arthropod detector for natural background scenes. The detector achieves an F1-score of 0.91, demonstrating strong performance despite substantial interspecific morphological variation and heterogeneity in photographic contexts.

We further demonstrated the taxonomical universality of the model showing high F1-score and IoU averaged at the class (0.79, 0.85) and order level (0.82, 0.86) and also a good detection generalizability (F1-score>0.90, IoU>0.83) on species, genera and families never encountered by the model during training. Finally, we show how this model can be improved to generalize to new datasets using data augmentation, complementary training data or fine-tuning and increase detection of small objects. In particular, we report performance of the improved models on three use cases largely used in non lethal insect monitoring: (i) diurnal pollinator monitoring through citizen science or (ii) flower and nocturnal insects monitoring through smartphone time-lapse of a UV-illuminated white panel.

These results mark an important step toward automated analysis of arthropod images in natural contexts, from both large-scale automated monitoring approaches or from citizen science monitoring programs.

## INTRODUCTION

Insects and other arthropods have been severely declining in recent decades (Hallmann et al., 2017; Seibold et al., 2019; van Klink et al., 2020). As insects play a key role in ecosystem functioning (Verma et al., 2023; L. H. Yang & Gratton, 2014), this could have many consequences, ranging from species extinctions (Kehoe et al., 2021) to a decline in pollination, impacting human food supply through lower agricultural yield (Tscharntke et al., 2012). This massive decline and its consequences calls for the development of better conservation and restoration strategies of insect populations (Kawahara et al., 2021; Miličić et al., 2021; Samways et al., 2020). Key to these efforts are efficient monitoring tools that can be deployed at a large scale are essential to quantify and highlight causes of decline and to evaluate restoration strategies (Krahner et al., 2025; Wagner, 2020).

Classical insect surveys are often limited in scale because they are time-consuming to deploy and require taxonomist or barcoding identification, thereby greatly restricting the number of sites that can be realistically monitored. Furthermore, most traditional methods rely on destructive sampling, such as pitfall traps, pan or malaise trapping (Montgomery et al., 2021), which raise ethical concerns, especially for species already in decline (Lövei & Ferrante, 2024). To address these challenges, there has been a recent push for non-destructive monitoring approaches (Barrett & Fischer, 2024), such as automated light traps to monitor moths (Bjerge et al., 2021; Jonason et al., 2014; Singh et al., 2022), pollinator camera traps on natural flowers (Darras et al., 2024; Naqvi et al., 2022) and artificial attractors (Sittinger et al., 2024), non lethal photographic traps (Chiavassa et al., 2024) and citizen science photographic surveys (Bedessem et al., 2022; Flaminio et al., 2021). While mitigating insect harm, these new arthropod monitoring approaches produce vast amounts of heterogeneous images that vary a lot between sensors and protocols used, and which are time consuming and difficult to process by hand (Alison et al., 2026).

To automate the insect monitoring process, computer vision methods have shown significant promise (Teixeira et al., 2023; Bjerge et al., 2021; Hartbauer, 2024; F. Yang et al., 2021; Alison et al., 2026; An et al., 2023; Grele & Richards, 2026). Large datasets of insect images labeled to the species, hosted on crowd-sourced platforms such as GBIF-mediated datasets, Observation.org and iNaturalist (Della Rocca et al., 2024; Horn et al., 2018; Robertson et al., 2019) have enabled the building of powerful insect classifiers for species recognition (Chiranjeevi et al., 2025; Roy et al., 2024; Stevens et al., 2024).

However, standard image classification methods suffer from several limitations, most notably their inability to handle images containing multiple arthropods despite such images being common in automated and citizen science monitoring tools (see Figure 1c). Such situations commonly arise when two individuals of the same species are present, leading to underestimation of insect abundance, or when multiple species co-occur in a single image (see Figure. 1d), leading to biases in assessment of presence and species richness. In contrast, object detection approaches in computer vision address this limitation by explicitly localizing objects of interest within an image (Diwan et al., 2023; Zhao et al., 2019). For each detected object, a detector typically outputs bounding box coordinates, thereby enabling the localisation, distinction and counting of multiple objects within the same image. Applying object detector algorithms to automated insect monitoring images could help resolve a major limitation of these approaches.

**Figure 1.**
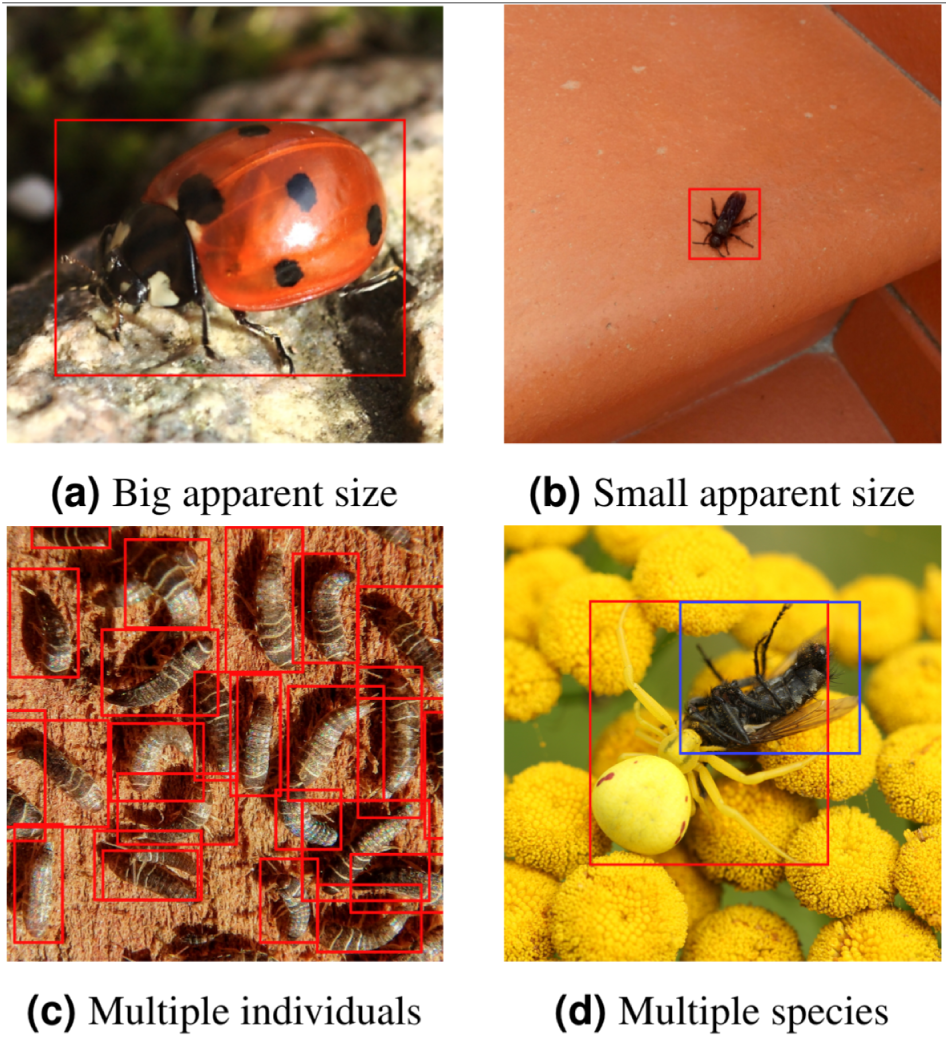
Visualization of arthropods labeled with their detection bounding box. Arthropods can have a big (a) or small (b) apparent size on the images. They also appear in varying numbers, with multiple individuals of the same species (c) or multiple species (d) on the same image.

Furthermore, modern deep learning based image classifiers have a maximum input resolution (Bakhtiarnia et al., 2024) due to hardware limitations, requiring resizing (most often to lower resolution) any image before using them as input for the model. Resizing to lower resolution can result in small objects, such as insects, being represented by only a few pixels (see Fig 1.b). Detecting insects in the original image first allows each insect to be cropped into a separate image. Since cropping happens before resizing to the classifier’s input resolution, the insect retains more pixels after resizing and has the potential to improve species identification. Detection-based cropping is now a common preprocessing step for classification of biodiversity images (Bjerge et al., 2025; Rigoudy et al., 2023). By limiting background information, object cropping can also help avoid shortcut learning, the tendency of deep neural networks to exploit spurious correlations instead of learning invariant, task-relevant representations, thereby impairing robustness and generalization (Geirhos et al., 2020; Rigoudy et al., 2023). The potential use of object detection for arthropods has already been highlighted (Stark et al., 2023; Teixeira et al., 2023) (Table 1), but application to realistic images remains rare.

**Table 1.** List of insect detection datasets, with public data available. The dataset size corresponds to the number of images and number of bounding boxes (in brackets). The number(#) of labels and Labels are the number of distinct classification labels and corresponding label names. The number of Classes, Order, Families, Genera and Species indicate the actual taxonomical diversity found in the source images, regardless of the bounding box labels. The background type corresponds to the type of environment the insects are photographed in.

| Source | Dataset name | Year | Dataset Size | # of Labels | Label names | # of Classes | # of Orders | # of Families | Background type |
| --- | --- | --- | --- | --- | --- | --- | --- | --- | --- |
| (Sun et al., 2018) | N/A | 2018 | 2,183 [729] | 6 | Bark Beetles species | 1 | 1 | 1 | uniform |
| (Chudzik et al., 2020) | GHCID | 2020 | 3,578 [4,406] | 1 | Grasshopper | 1 | 1 | N/A | natural |
| (Wang et al., 2020) | Pest24 | 2020 | 25,378 [192,408] | 24 | Coleoptera, Homoptera, Hemiptera, Orthoptera, Lepidoptera | 1 | 4 | 7 | uniform grid |
| (Drange, 2026) | ArTaxOr | 2020 | 15,376 [19,605] | 7 | Araneae, Coleoptera, Diptera, Hemiptera, Hymenoptera, Lepidoptera, Odonata | 2 | 7 | N/A | natural |
| (Supervisely, 2022) | Bee Image Object Detection | 2022 | 3,044 [21,240] | 1 | Bee | 1 | 1 | 1 | hive entrance |
| (Bjerge et al., 2025) | N/A | 2025 | 777 [5,817] | 1 | Insect | 2 | 15 | N/A | uniform |
| (Svenning et al., 2025) | flatbug | 2025 | 6,131 [113,552] | 1 | Arthropod | $\geq 2$ | $\geq 7$ | N/A | uniform / natural |
| (Ştefan et al., 2025) | OOD dataset | 2025 | 23,899 [24,838] | 6 | Hymenoptera, Diptera, Coleoptera, Formicidae, Araneae, Hemiptera | 2 | 5 | N/A | natural |
| **ours** | **ArthroNat** | 2025 | 13,661 [19,663] | 1 | Arthropod | **11** | **67** | **749** | natural |

Many insect detectors trained on specific datasets suffer from the out-of-distribution (OOD) problem, whereby detection performance degrades when the model is applied to images whose statistical properties (e.g., background structure, species appearance, taxonomical range) differ from those encountered during training (Ștefan et al., 2025). Indeed, most datasets publicly available tend to focus on specific use cases, in terms of image background and target taxon (Stark et al., 2023; Svenning et al., 2025). Insects are often presented on uniform backgrounds (Bjerge et al., 2025; Sun et al., 2018; Q.-J. Wang et al., 2020), and when this is not the case, only a handful of taxa are covered (Chudzik et al., 2020; Ștefan et al., 2025). In non-destructive arthropod monitoring applications, such distribution shifts may arise from changes in sampling approach (passive camera trap vs active citizen photographic acquisition choices), taxa monitored (soil arthropods vs diurnal pollinators vs moth), sampling technology (camera type or resolution), photographic background and illumination linked to sampling technology (homogenous background and artificial light automated light moth trap and non lethal photographic trap vs natural background and light in citizen survey or pollinator camera trap on flowers) or post sampling image processing (manual crop and/or choice of best photography for identification in citizen science protocols). Indeed, the most comprehensive collection of annotated arthropod images for detection and segmentation compiled recently (Svenning et al., 2025, Table 1) is likely to be useful for only a subset of species and monitoring systems. This is because the dataset is largely dominated by specimens photographed against flat and relatively uniform backgrounds taken with standardized sensors, which limits model generalization when applied to live individuals in natural environments (Mazen, 2023; Svenning et al., 2025).

We aim to provide a general detection and classification tool by simultaneously building a dataset and training a model for detecting any order, family, genus or species of live arthropods in many natural backgrounds in which they occur during citizen science or automated non destructive monitoring field protocols. We present our methodology for systematically gathering, quickly labeling, and validating a dataset that encompasses a wide diversity of terrestrial arthropods, ensuring both annotation quality and taxonomic representativeness. Building on this foundation, we perform an extensive analysis of model generalization, assessing performance not only on unseen data within the dataset but also on external datasets as well as on novel species, genera, and families. In doing so, we address 4 specific research questions:

1: How many images and iterative annotation steps are necessary to build a robust (F1-score > 0.9) detector?
2: How generalist is the detector to biological taxonomy (generalization within Class and Order and between Family, Genus and Species)?
3: How can the detector be improved for out-of-distribution datasets?
4: How does the model perform in real word applications of non destructive insect monitoring?

## MATERIAL & METHODS

### Training Data

To construct a database for training our detection algorithm, we used the TAXREF database (the official taxonomic reference for France (Gargominy et al., 2020) to compile a list of terrestrial arthropod species present in metropolitan France. Among the 47,299 species recorded in TAXREF, 32,154 were also present on iNaturalist, which corresponded to 769 families (out of the 1081 families existing according to TAXREF). However, 20 of those families had no observations on iNaturalist, meaning we only had **749** arthropod families in reality.

We downloaded images of arthropod species from the iNaturalist platform in October 2024 (Figure 1). Only observations labeled as “Research Grade” were included, indicating that the species identification was confirmed by the community (at least 2/3 of identifications agree on the taxon). Within each family, the species with the greatest number of iNaturalist observations was selected for the annotation phase. We wanted to maximize coverage by representing all the families, while making sure that the model would perform well on the most likely-to-be-encountered species. We used only one image per iNaturalist observation to maximize image diversity and avoid having near-duplicates in the dataset. The pictures depicted in Figure 1 constitute an example of such collected images, while Figure 2 illustrates all families, orders and classes included in the dataset. All images of the training dataset can be downloaded using our dedicated script on GitHub (github.com/edgaremy/arthropod-detection-dataset).

**Figure 2.**
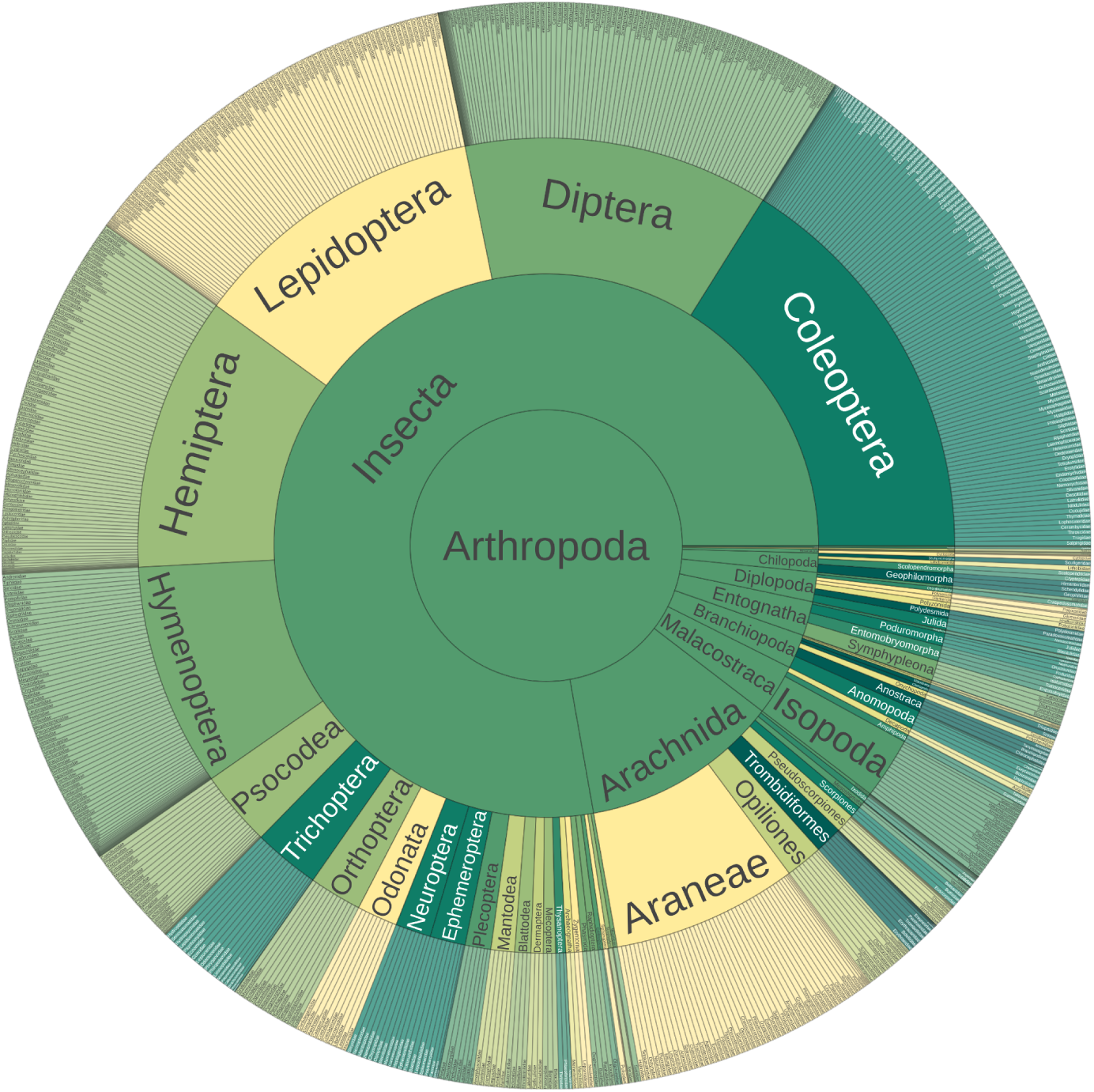
Taxonomic visualization of the data contained in the ArthroNat detection dataset. All arthropod families are on the exterior of the circle. Going towards the center of the circle means going to the parent taxon, up in the taxonomic tree. The levels depicted from the outside to the inside: family, order, class, and the phylum of interest, *Arthropoda*. The size of each section is proportional to its respective number of images (the color variations are only here for cosmetic and readability purposes).

### Iterative Data Annotation and Model Training

Since manually drawing bounding boxes around the objects is a tedious and time-consuming task, we adopted a semi-automated approach in which human annotators only corrected model predicted annotations. We chose an iterative approach (Adhikari & Huttunen, 2021) that combined image annotation and model training with the objective of obtaining a final detector that reached a 90% F1-score while minimizing human annotation effort.

The iterative data annotation process consisted in repeating the following steps. We first collected a batch of images, taken from our curated data listing (one image for the most represented species of each of the 749 families). We then pre-annotated the data using a YOLO11 nano (Ultralytics, 2024) model trained on the previously annotated batches (except for the first batch, for which a model pre-trained on COCO (Lin et al., 2015) was used). We presented the model predictions to human annotators, who validated or corrected the bounding boxes on each image of the batch using a custom made interface. Depending on the cases, this could mean slightly adjusting the borders of the box, or going as far as removing an erroneous detection or adding a missing box. In many cases however, the model’s pre-annotation required little to no correction in order to obtain an accurate label. This allowed for bounding box labels to be produced at a much faster pace than if done entirely by hand. We processed a total of 22 consecutive batches, making for a total of 13,661 images. Note that some rare families had less than 22 observations in total available on iNaturalist, so the batch size slightly decreased from one batch to the next because of them.

We used the YOLO11n model as it is the smallest in the YOLO11 family (see Ultralytics, 2024), allowing for faster training, in order to quickly iterate through the annotation process. Every model training in this paper was done using one or two Nvidia Quadro RTX 5000 GPUs, with training time ranging from a few hours up to around one day, depending on dataset size, model size and number of GPUs used. For each training, the number of epochs was set to 100, 90% of images were used for training, while the remaining 10% were kept for validation, every other parameter was left at its default value.

The model’s performance was measured using the IoU (Intersection over Union) and the F1-score. The IoU is computed by comparing the expected bounding box label against the bounding box predicted by the model: it is the ratio of the intersection of their respective areas over the union, and is a metric designed to assess the quality of a bounding box. If no bounding box is associated with a label in the detection process, the corresponding IoU was set to zero. The F1-score (as well as precision and recall) depends on the IoU threshold, as this threshold sets what is considered a Positive or a Negative detection. We set the IoU threshold as well as the model confidence threshold to 0.5. This can be considered standard practice for assessing a detection model’s performance, and simplifies the task of comparing multiple models (Salari et al., 2022). F1-score and IoU were the main metrics used throughout our analysis, as F1-score encapsulates both precision (what percentage of detections are actual arthropods) and recall (what percentage of arthropods are actually detected), illustrating the trade-off between the two with a unique score. The IoU metric could be assessed for each bounding box individually, however, the F1-score was computed at the image level.

We first validated our iterative training process, by retrospectively measuring the model’s performance at each stage of the process. More specifically, as our iterative annotation process added new training data per batch, once the entire data was properly labeled, we could emulate other training scenarios in which each annotation batch arrived in a different order. The objective was to make sure that performance growth after each batch was not due to chance, but in fact was reproducible regardless of the batch order chosen. The results were validated, using a 5-fold cross-validation. Using the 22 annotated image batches at our disposal, we simulated alternative scenarios, in which the batches would have arrived in a different order. For the first batch, we drew one random batch for training and one random batch for validation, out of the 22 at our disposal. The 20 remaining batches are used for testing. This was repeated 5 times to obtain 5 distinct scraping/training scenarios. For the second batch, we drew two random batches for training, one other for validation, and the rest for testing, and repeated this 5 times. By doing this, we continued up until batch number 20, in order to have 5 random scenarios for each step of the iterative scraping process. Note that we stopped at 20 in order to use the two random remaining batches for validation and testing. For each scenario, a YOLO11n model was trained, starting from the same model weights we used in the actual iterative process.

To build the final detection model (baseline model) and associated dataset, called **ArthroNat**, all images used in the iterative training were randomly split per species into training, validation, and test sets, targeting an 80/10/10 ratio while ensuring at least one image per split. For species with fewer than three images, if there was only one image, it went in the training set, if there were only two images, one went in the training set and the other to the testing set. Applying this rule resulted in an overall distribution of 71.4%/13.7%/14.9%. A YOLO11l (large), which presented the best compromise between performance gains and computational cost (see also Figures S10 and S11 in Supplementary) was then trained on 100 epochs.

### Generalization across arthropod taxonomy

We assessed taxonomic generalization of our baseline model using two main approaches. Both approaches used F1-score and IoU metrics, allowing us to quantify the model’s potential as a general-purpose arthropod detector. First, we measured detection performance by taxonomic class and order. Second, we tested how well the model generalized to new species: We defined four levels of taxonomic generalization (see Table 2):

1. New images of the same species as those in the training set (the ArthroNat test set).
2. Images of the same genus as the training set, but different species, using the second most documented species of each family.
3. Images from the same families as the training set, but different genus.
4. Images from arthropod families not present in our dataset, sourced from a different geographical region.

**Table 2.** Summary of the different taxon levels of generalization used to assess the model’s performance on other taxa.

| Level of generalization | Description | # of images |
| --- | --- | --- |
| 1. Same species | *Test set from ArthroNat (our dataset)* | 2,037 |
| 2. Same genus (diff. species) | A different species for each family in our data | 361 |
| 3. Same family (diff. genus) | A different genus for each family in our data | 435 |
| 4. Diff. families | Arthropod families not present in our data (South American) | 215 |

For the across-family generalization assessment, we collected images of arthropods observed in South America from iNaturalist, using the criteria −55 < latitude < 15 and −85 < longitude < −30. We excluded any arthropods belonging to families present in our dataset and manually removed marine species using the WoRMS platform (WoRMS Editorial Board, 2025).

Images for the first level are equivalent to the ArthroNat test split, while data from level 2 to 4 was independently collected from iNaturalist and manually annotated. Table 2 summarizes each taxon level of generalization, indicating also the number of images.

### Generalization to new datasets

Beyond taxonomic generalization, the ability to generalize to new datasets compiling images acquired using different imaging systems and under different environmental conditions is a fundamental requirement for a general detector of arthropods. To illustrate how our detector could be adapted to real world applications, we gathered 3 independent test datasets (Figure 3) that represented current practice in non lethal arthropod monitoring:

– The SPIPOLL dataset presents a wide diversity of pollinator images, including 548 insect taxa observed on 473 plant species, collected by 256 participants in a French citizen science program (*Spipoll – Suivi photographique des insectes pollinisateurs*, 2026) dedicated to monitoring plant–pollinator interactions (see SI for more details on the program and image selection).
– The OOD dataset (“*Dataset of arthropod flower visits captured via smartphone time-lapse photography*”, also presented as an Out-Of-Distribution dataset (Ștefan et al., 2025)) contains various timelapse images shot on a smartphone using a tripod over open flowers. This dataset depicts the kind of data that can be obtained from an automated smartphone based-pollinator monitoring program, and its data is publicly available.
– The LEPINOC dataset is composed of data produced by Lépinoc, a citizen science program for monitoring moths, conceived and led by the French association Noé (*Association pour la biodiversité – Noe.org*, 2026). It also contains timelapse images taken on a smartphone using a tripod, but this time pointed at a white UV-illuminated sheet during the night. High-resolution (3072 × 4096 pixels) Lepinoc images were tiled into 24 640 × 640 patches to maximise object apparent size. Note that every training, validation and statistics using Lepinoc is referring to the dataset with the 640 × 640 tiled images.

**Figure 3:**
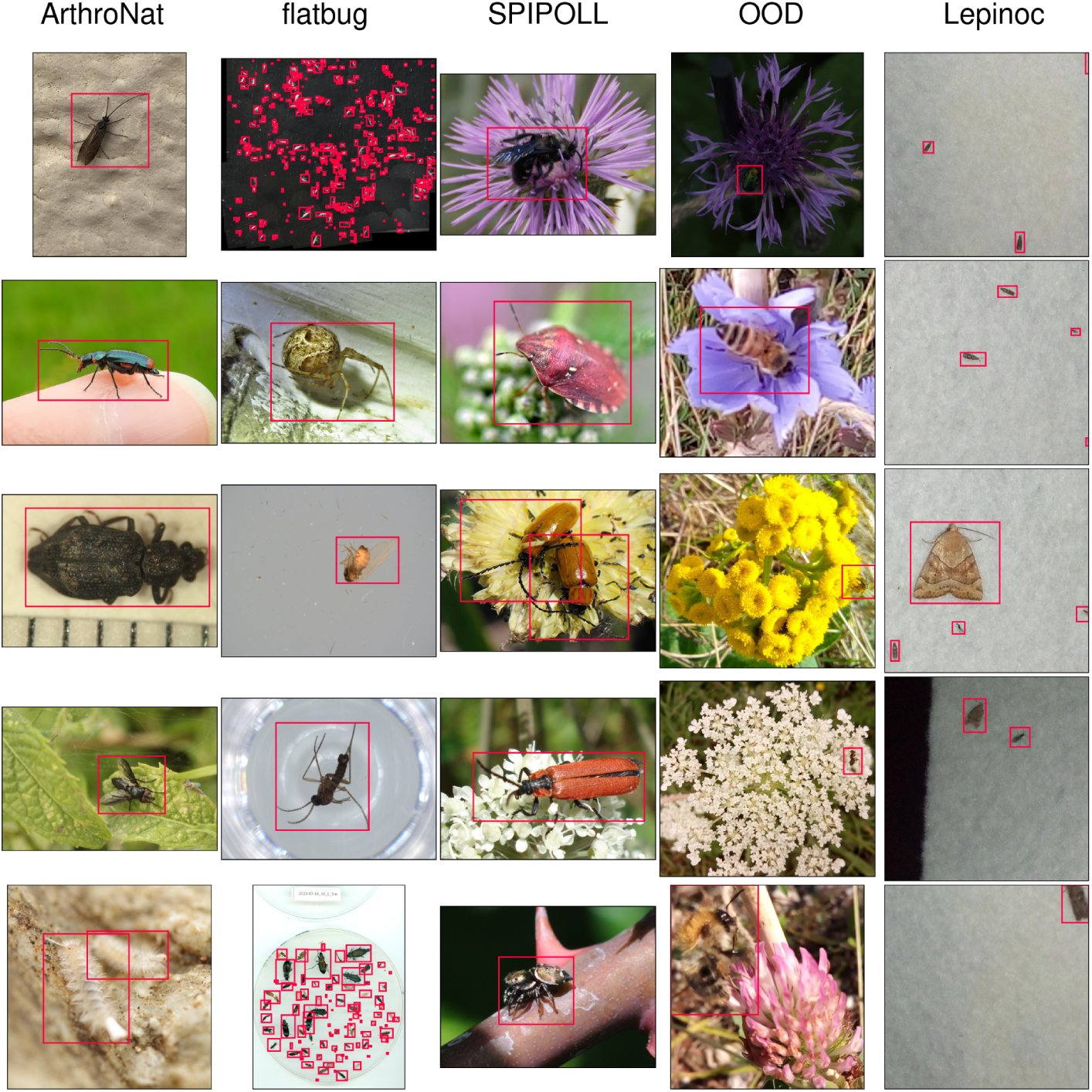
Examples of images from each generalization test set, with bounding box label visualization in red.

Image properties of ArthroNat and SPIPOLL dataset were largely shared, with one or a small number of relatively large arthropods on natural backgrounds, captured by human observers. In contrast, the smartphone OOD and LEPINOC datasets largely differed from our initial dataset by the apparent size of the arthropod in the picture and the fact that they were taken by a fixed smartphone. We defined apparent size as the area of an arthropod bounding box relative to the total image area (see Figure 1.a and 1.b). Note that it is determined mainly by factors such as the use of a macro lens and the distance to the subject, rather than the insect’s actual size. In addition, the LEPINOC dataset differed from ArthroNat by the white background, nocturnal imaging conditions and higher number of individuals per image (Table 3).

**Table 3:** Specifications of each generalization test set. The number (#) of test images indicates the total number of images available for testing (this only includes the test splits for ArthroNat and flatbug, as the rest of data was used for training). The number of bounding boxes (# bboxes) corresponds to the total count of bounding boxes contained in the test images.

| Dataset | # test images | # bboxes | bbox apparent size (% of image) |  |  |  | # bbox per image |  |  |  |
| --- | --- | --- | --- | --- | --- | --- | --- | --- | --- | --- |
|  |  |  | mean | std | min | max | mean | std | min | max |
| ArthroNat | 2,037 | 2,812 | 0.25 | 0.25 | $3.4e-4$ | 1.00 | 1.38 | 2.07 | 1 | 42 |
| flatbug | 1,139 | 21,825 | 0.01 | 0.05 | $2.9e-6$ | 0.87 | 19.16 | 50.64 | 0 | 577 |
| SPIPOLL | 1,857 | 1,998 | 0.37 | 0.19 | $9.9e-4$ | 0.92 | 1.08 | 0.43 | 1 | 10 |
| OOD | 2,391 | 2,498 | 0.07 | 0.09 | $1.1e-3$ | 0.47 | 1.04 | 0.24 | 1 | 3 |
| Lepinoc | 770 | 1,525 | 0.02 | 0.04 | $9.1e-6$ | 0.63 | 1.98 | 1.69 | 1 | 13 |

We then proposed and evaluated three dataset generalization strategies (DGS) that differed in annotation effort and computational cost, depending on the target application: (DGS1) using data-augmentation methods during training specifically designed to improve adaptation to images containing multiple small objects, (DGS2) adding complementary training data and (DGS3) fine-tuning our model using a limited amount of target-application data.

In DGS1, we used mosaicing, an data augmentation method that creates a grid (mosaic) comprised of multiple images, generally 2×2 or 3×3 images. This artificially increases the number of insects per image and reduces their apparent size—mimicking the properties of datasets like flatbug, which naturally have more and smaller insects per image. We implemented 4 mosaicing scenarios (none, 2×2, 3×3, 4×4) to see how well this approach worked. Note that the Ultralytics library enables 2×2 mosaic augmentation by default (see Figure S12 and the dedicated section in the Supplementary Information for details), reflecting common practice in object detection, including insect detection (Mazen, 2023).

For the DGS2 we used the recently released flatbug dataset (Svenning et al., 2025) as a complementary training data that presents images with a high number of small (often dead) annotated arthropods per image (Figure 3, Table 3). Flatbug annotation was first converted from segmentation to detection format. We then trained two new detection models, one using only flatbug data (flatbug) as a control and one combining our initial ArthroNat dataset with the flatbug dataset (ArthroNat + flatbug). The ArthroNat+flatbug dataset contains in its train/validation/test split respectively 13,972 (9,051 + 4,921), 1,863 (1,737 + 126) and 3,044 (1,905 + 1,139) images. Unless otherwise stated, results were reported for YOLO11l, with additional YOLO11n comparisons in the Supplementary Information. Performance for DGS1 and DGS2 was evaluated on five test sets (Table 3): ArthroNat test set, flatbug test set and the three application datasets SPIPOLL, OOD, and Lepinoc. To evaluate how DGS1 and DGS2 improve performance according to object apparent size, we computed F1-score and IoU by size decile of the ArthroNat test. Note that the deciles are not the same for F1-score and IoU, as F1-score is computed per image, while IoU is computed per bounding box.

For our third generalization approach (SDG3), we evaluated whether targeted fine-tuning of the best-performing model from SDG1 and SDG2 (ArthroNat+flatbug) could improve transfer to specific applications (OOD and Lepinoc) with limited annotation effort. Fine-tuning means starting from an already trained model and adapting it to a new dataset with a small amount of additional training data. Because this is a final adjustment step, training is typically shorter and more conservative so that the model adapts to the new context without drifting too far from previously learned representations (Chu et al., 2016; Church et al., 2021; Poojary & Pai, 2019; Yin et al., 2017). To implement this, both application datasets were split into train/validation/test partitions (80%/10%/10%). The 80% train and 10% validation splits were both used during fine-tuning: the validation split was used to monitor validation performance and select the best checkpoint. Only the post-split 10% test partitions were used for the evaluations reported. For OOD, this split was performed temporally, assigning the oldest data to the training set and the most recent to the test set, while for Lepinoc, we ensured that all tiles originating from the same image were assigned to the same split.

Using the 80% training partitions, we then constructed fine-tuning subsets of increasing sizes (100, 500, 1,000, and 2,000 images) each comprising five folds. Each subset was designed to preserve the diversity of the original data, spanning all available dates for OOD and all tiled images for Lepinoc. Validation subsets were set to 20% of each training subset size, and the held-out test split remained unchanged for all evaluations. We reported 5-fold fine-tuning performance using F1-score and IoU, separately fine-tuned and tested once on OOD and once on Lepinoc. Fine-tuning was conducted using default parameters of Ultralytics, with the following exceptions: 20 epochs with no warmup, AdamW optimizer with a starting learning rate of 0.001 and final learning rate of 1%, and the 10 first layers of the model frozen, which effectively means that we don’t modify the backbone of the model, and only fine-tune its head. Those fine-tuning settings were selected after trying a few cases. For comparison, we also performed a control fine-tuning experiment with the same procedure, but using the pre-trained COCO weights provided by Ultralytics as starting point.

## RESULTS

### Iterative training

During the iterative training, we processed a total of 22 consecutive batches. Making for a total of 13,661 images (9,754 for training, 1,870 for validation and 2,037 for testing) with at least one bounding box. 212 images showing no actual arthropods, but only traces (e.g., galls), had no bounding boxes and were therefore excluded. In total, the data covers 11 classes, including 67 orders, amounting to 749 arthropod families (Figure 2) (out of the 1081 existing French terrestrial arthropods). The dataset image distribution by taxonomic level is as follows: 1,235 images per class on average (*std* = 3,128, *min* = 6, *max* = 10,565), 203 images per order on average (*std* = 439, *min* = 2, *max* = 2,187) and 18 images per family on average (*std* = 6.6, *min* = 2, *max* = 22).

The iterative training shows an overall increase of detection performance with each image batch added to the training, for every evaluation metric (F1-score, Precision, Recall, Mean IoU, Figure 4). After the first batch, the model already reached a mean F1-score of 0.77 (*std* = 0.01, *best* = 0.80, *worst* = 0.75) and an average IoU of 0.69 (*std* = 0.01, *best* = 0.71, *worst* = 0.65). The increase is more noticeable within the early batch additions (up until around 2,000 images), and after that, the performance seems to reach a plateau, or at least increase more slowly. At the twentieth and last batch addition (13,661 images), an average F1-score of 0.9 is reached (*std* = 0.02, *best* = 0.91, *worst* = 0.87), illustrating both good precision (*mean* = 0.91, *std* = 0.01, *best* = 0.93, *worst* = 0.90) and recall (*mean* = 0.90, *std* = 0.02, *best* = 0.92, *worst* = 0.87). At the same time, an average IoU of 0*.8*3 is reached (*std* = 0.02, *best* = 0.84, *worst* = 0.80).

**Figure 4.**
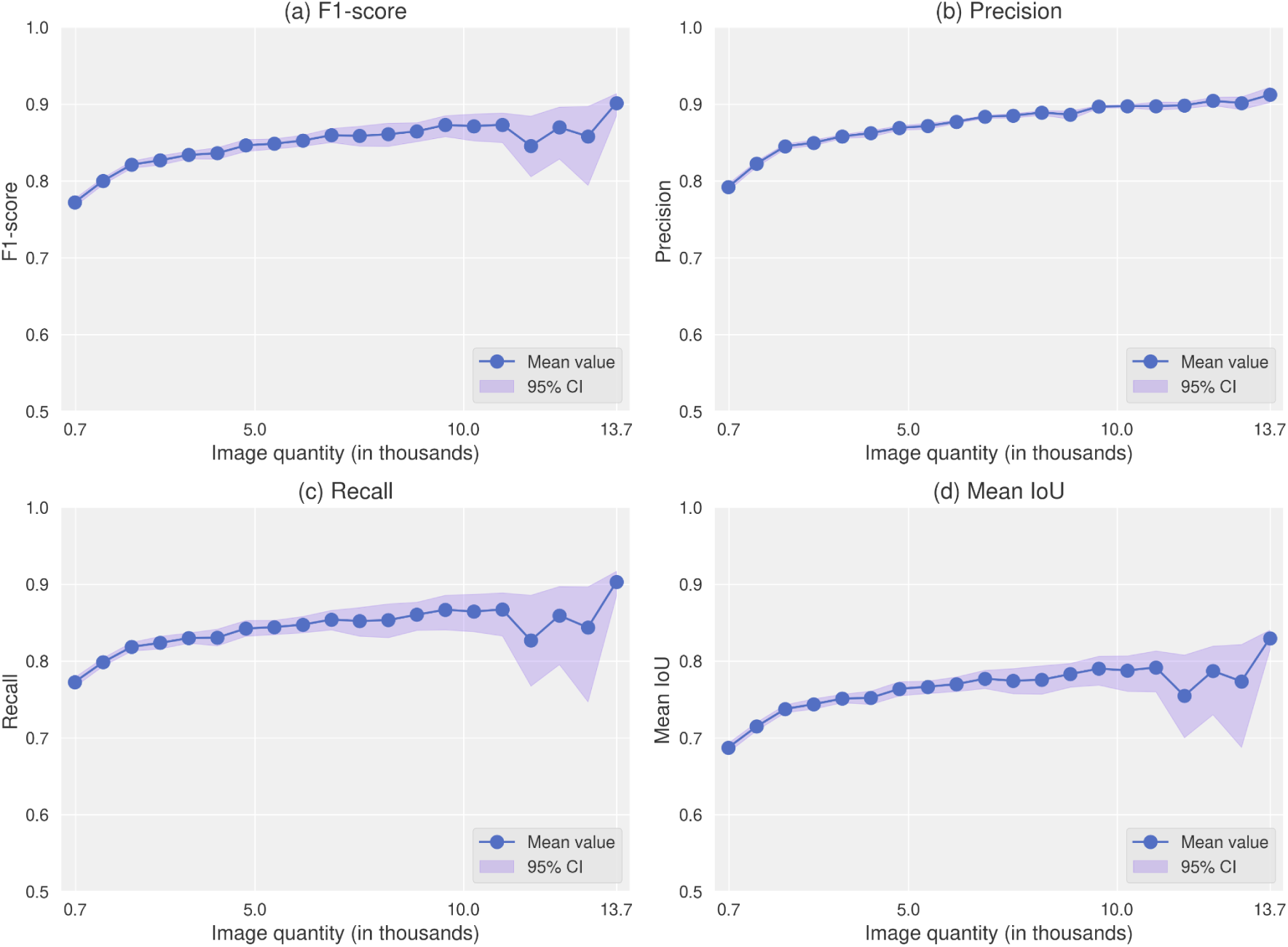
Cross-validation of the detection performance of the models, obtained throughout the iterative annotation process. For F1-score (a), Precision (b), Recall (c), and mean IoU (d), points represent the mean for a given image quantity, and the shaded area represents the 95% confidence interval.

#### Performance based on taxonomy

In order to identify any discrepancies in detection performance between different arthropod taxa, we evaluate the F1-score and mean IoU (Table 4) of our baseline model on each taxonomic class of the ArthroNat test set. For a finer analysis, we also evaluate the same metrics averaged by taxonomic order (Table 5). The F1-score is averaged across images, while the IoU is averaged across bounding boxes. Because the number of images per taxon varies widely, metric estimates for classes or orders with few images can be less reliable. Also note that taxon-level averages treat each class/order equally, so rare groups can have a disproportionately large effect on these averages compared with image-level metrics, which weight each image equally.

**Table 4.**
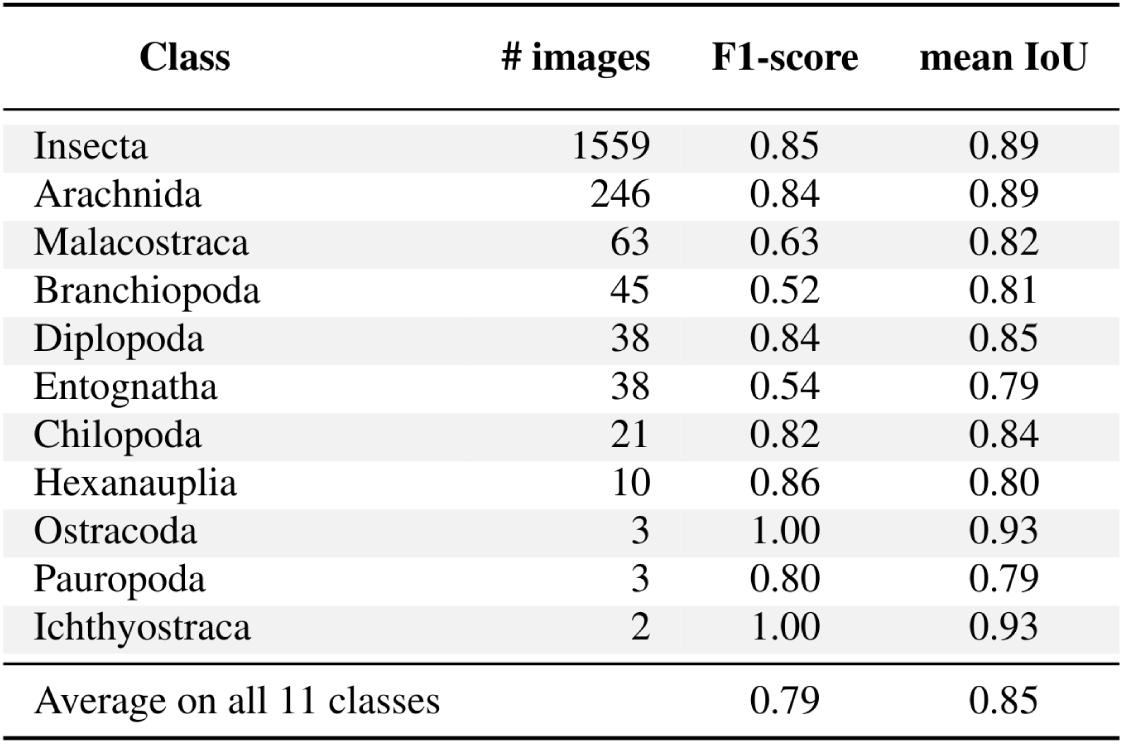
Class-level F1-scores and mean IoU of our baseline model on the ArthroNat test set.

**Table 5.** Order-level F1-scores and mean IoU of our baseline model on the ArthroNat test set (top 20 with most images out of 67, see Table S2 for the complete table).

| Order | # images | F1-score | mean IoU |
| --- | --- | --- | --- |
| Coleoptera | 315 | 0.92 | 0.92 |
| Diptera | 251 | 0.86 | 0.92 |
| Lepidoptera | 220 | 0.94 | 0.91 |
| Hemiptera | 212 | 0.76 | 0.87 |
| Hymenoptera | 190 | 0.83 | 0.90 |
| Araneae | 144 | 0.91 | 0.92 |
| Psocodea | 64 | 0.78 | 0.86 |
| Trichoptera | 59 | 0.88 | 0.86 |
| Isopoda | 50 | 0.59 | 0.81 |
| Orthoptera | 38 | 0.87 | 0.84 |
| Ephemeroptera | 32 | 0.82 | 0.86 |
| Opiliones | 30 | 0.85 | 0.87 |
| Odonata | 29 | 0.98 | 0.90 |
| Neuroptera | 26 | 0.96 | 0.91 |
| Trombidiformes | 24 | 0.66 | 0.78 |
| Plecoptera | 23 | 0.81 | 0.93 |
| Pseudoscorpiones | 21 | 0.85 | 0.89 |
| Mantodea | 18 | 0.74 | 0.76 |
| Anomopoda | 17 | 0.83 | 0.88 |
| Symphyleona | 14 | 0.67 | 0.90 |
| Average on all 67 orders |  | 0.82 | 0.86 |

We tested whether the number of images per taxon was related to detection performance using simple linear regression models (number of images as a function of ArthroNat performance metric), at both class and order levels, for F1-score and mean IoU. In all four models, the regression slopes were not significantly different from zero (p-values between 0.184 and 0.781), and the relationships were weak (R² values between 0.005 and 0.074). Overall, these analyses indicate no significant linear association between number of images by taxon and model performance on the ArthroNat test set.

The mean IoU maintains similar quality of bounding boxes across taxa, for both class– and order levels, averaging at 0.85 or above. However, for both class– and order levels, the F1-score average is slightly degraded for some taxa such as the Branchiopoda and Entognatha classes, and the Isopoda order, which all have F1-scores below 0.6.

To assess how well our model generalizes to increasingly distant taxa, we also compare its performance (F1-score, Precision, Recall and IoU) across the four generalization levels defined in Table 2: Same species, Same genus but different species, Same family but different genus, and Different families. The performance on new taxa is close to (if not better than) the performance on taxa present in the training set (“*same species”* level) with F1-score above 0.9 and IoU above 0.8 (Figure 5). Note that the performance on the “*same species”* level (blue bars in the figures) is strictly equivalent to the overall performance reported in Table 6, as the ArthroNat test set is composed of the same species as the training set. It is also expected that performance on the ArthroNat test set differs from the performance at the end of iterative training (Figure 4), because the final ArthroNat split is not identical to the splits made during cross-validation.

**Figure 5.**
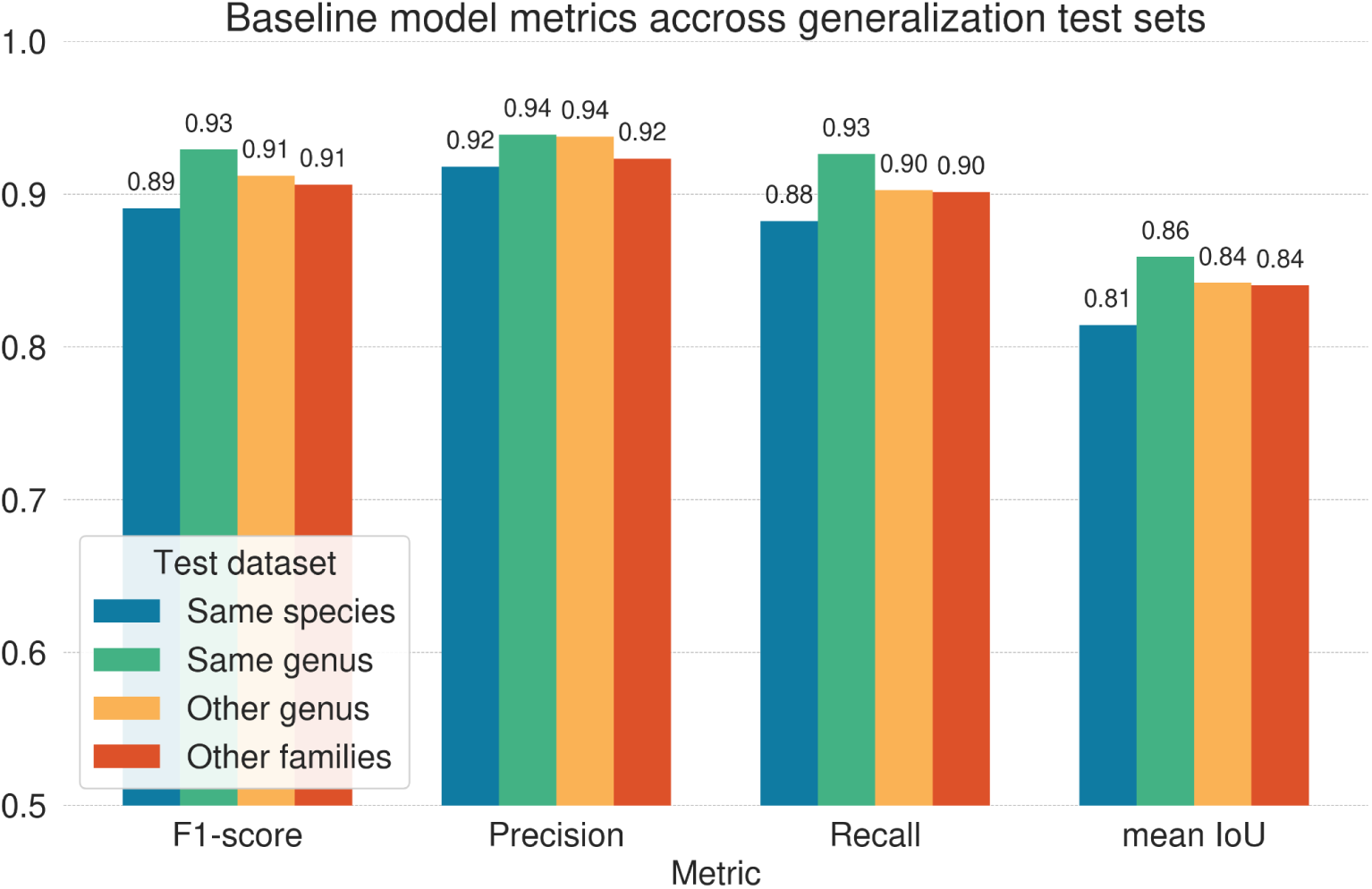
Comparison of the average F1-score, Precision, Recall and mean IoU of our baseline model across the different taxon generalization levels. The various taxon levels are described in Table 2. The mean F1-score, Precision and Recall are computed by averaging the metric of each image in the set. The mean IoU is computed by averaging the metric of each image in the set.

**Table 6.**
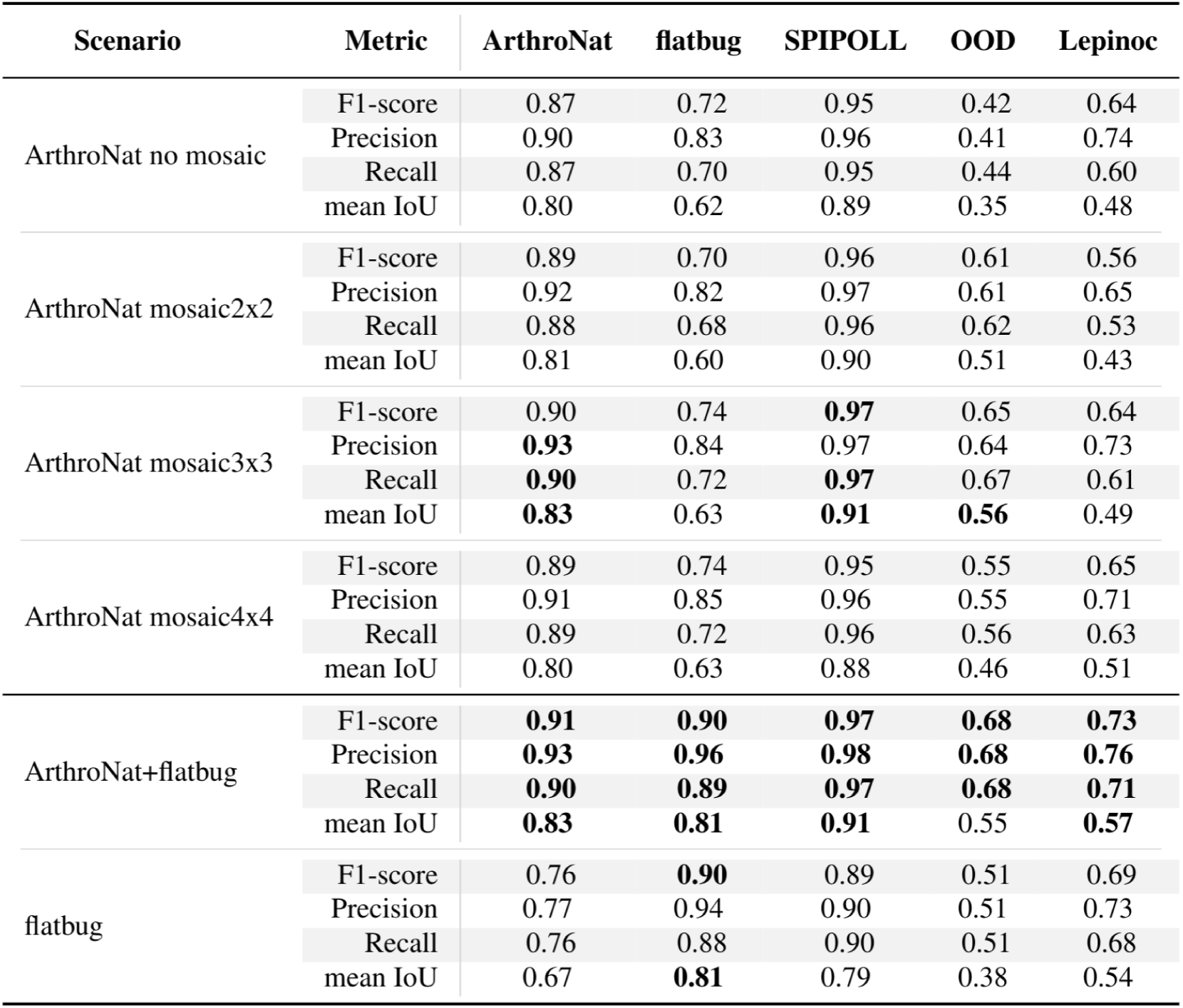
Comparison of the model’s average performance (F1-score, Precision, Recall and mean IoU) on the ArthroNat, flatbug, OOD, Lepinoc and SPIPOLL test sets, depending on data augmentation method and training datasets. The mean IoU is computed for each image and then averaged. Best values for a given (metric, dataset) pair are highlighted in bold.

### Generalization to new datasets

Regardless of the data generalization strategy (DGS) used, detection performance remains consistently higher on the ArthroNat and SPIPOLL test sets, with very high scores reaching up to 0.97 in F1-score and 0.91 in IoU for the latter citizen science dataset (Table 6). We also observe that detection performance increases with arthropod apparent size deciles, reaching a plateau for objects occupying 30-40% of the image (Figure 6). Notably, good performance (F1-score > 0.8, IoU > 0.7) is already achieved for arthropods representing as little as 5-7% of the image.

**Figure 6.**
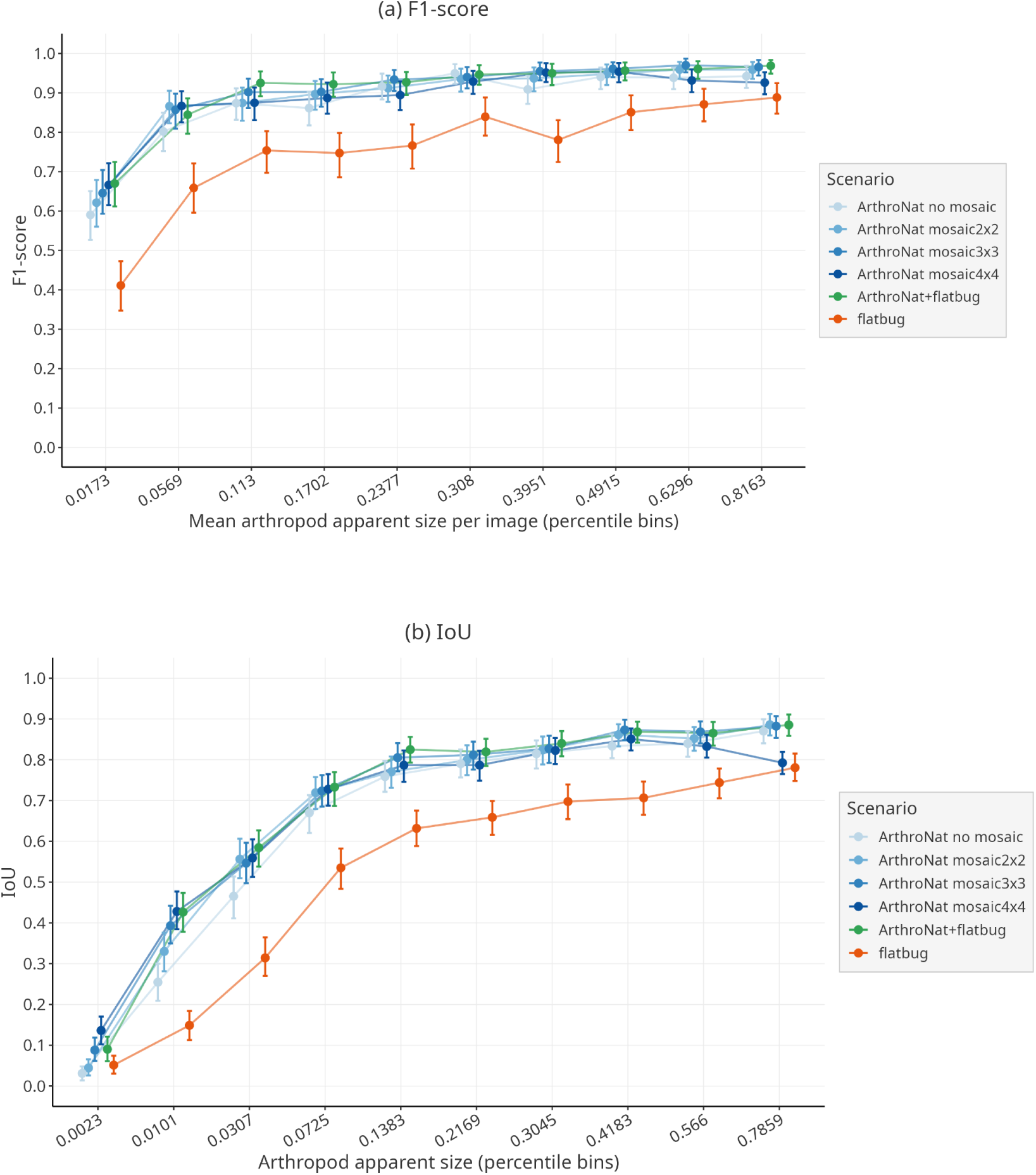
F1-score and IoU on the ArthroNat test set, depending on arthropod apparent size (grouped by decile), compared across our training scenarios. Mean and 95% confidence intervals are represented for F1-score (a), according to decile average apparent size per image, and for IoU (b), according to decile average apparent size across bounding boxes.

We then examine whether mosaic-based data augmentation during training could help generalize to new datasets (DGS1) and in particular whether increasing the level of mosaicing enhances detection on small objects and thus performance on dataset with smaller average arthropod apparent size (flatbug, OOD, LEPINOC; Table 3).

On the ArthroNat test set, detection performance appears to increase approximately linearly with mosaicing increase for the smallest object sizes, specifically within the first decile for the F1-score and the first two deciles for the IoU (Figure 6). For higher deciles (i.e., larger object sizes), 3×3 mosaicing consistently yields the best or among the best performance in terms of both F1-score and IoU (Figure 6). In contrast, training without mosaicing generally results in the lowest or among the lowest performance across all size deciles. We also observe a trade-off with 4×4 mosaicing: while it outperforms lower-level mosaicing for the smallest objects (first decile), its performance decreases for larger object sizes (last decile), particularly in terms of IoU.

At the dataset level, training with 3×3 mosaics data augmentation although seems to give best detection performance except for generalization on flatbug and LEPINOC dataset (which are the two datasets with the smallest arthropod average apparent size) for which 4×4 mosaics training give respectively similar or slightly higher performance (Table 6).

In DGS2 we evaluate how integrating the flatbug dataset to the training helps to generalize to other datasets. We found that this strategy gives equivalent results to 3×3 mosaic data augmentation on test sets with large average arthropod apparent size and photographs taken by citizens (ArthroNat and SPIPOLL) but outperforms any mosaic training for other dataset with smaller arthropod apparent size (flatbug, LEPINOC, OOD; Table 6). We also show that combining ArthroNat and flatbug datasets during training (Arthronat+Flabug model) gives better results on all test datasets than the a equivalent detector trained only on flatbug data except for the flatbug test set for which detection performances are equivalent with both models.

Overall the Arthronat+Flabug model achieves a high F1-score and mean IoU of respectively, 0.91 and 0.83 on ArthroNat, 0.90 and 0.81 on flatbug, and 0.97 and 0.91 on SPIPOLL. Detection performance is lower on smartphone based datasets with F1-score and mean IoU of 0.68 and 0.55 on OOD, and 0.73 and 0.57 on Lepinoc.

In DGS3, we thus fine-tune our best performing model (the ArthroNat+flatbug scenario) on the two testing sets where it struggles the most; the OOD test set (Figure 7) and the Lepinoc test set (Figure 8). In both cases, with the same number of target-domain images, our model consistently achieves better detection performance (F1-score and IoU) than the control model (pre-trained on COCO). Using 2,000 images for fine-tuning, our model improves its F1-score and mean IoU by about 5 percentage points on average. On the OOD test set, F1-score increased from 0.68 to 0.74 and mean IoU from 0.55 to 0.62. On the Lepinoc test set, F1-score increased from 0.73 to 0.78 and mean IoU from 0.57 to 0.61. For both datasets, fine-tuning with 100 images did not yield measurable improvements. On Lepinoc, improvements were already observed with 500 images (F1-score from 0.73 to 0.76, mean IoU from 0.57 to 0.60 on average), whereas on OOD, similar gains emerged after 1,000 images (F1-score from 0.68 to 0.71, mean IoU from 0.55 to 0.60).

**Figure 7.**
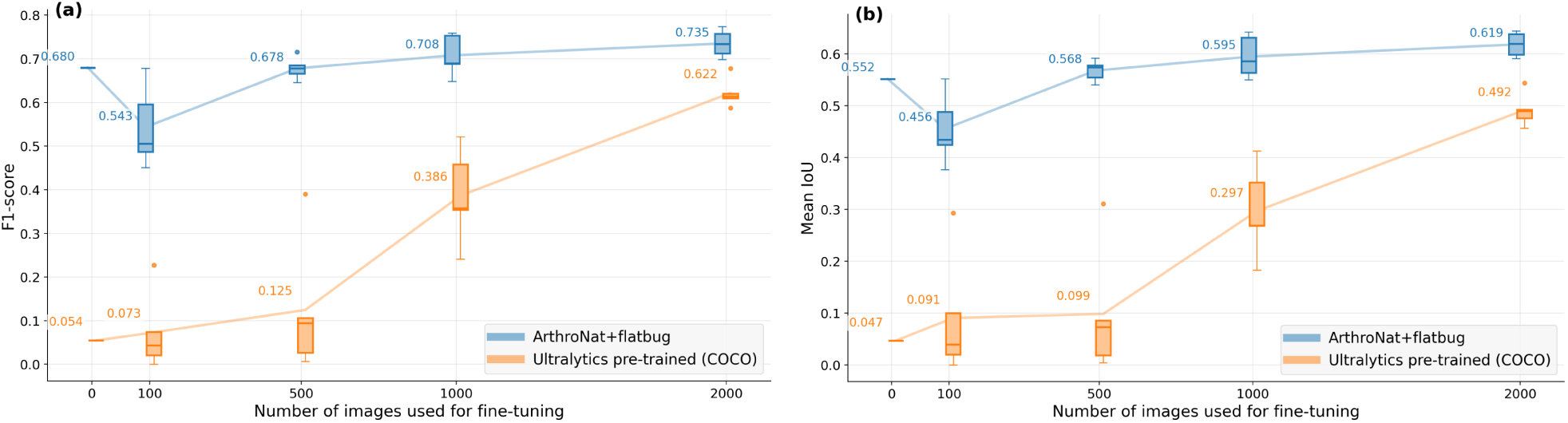
F1-score (a) and mean IoU (b) as a function of the number of images used for fine-tuning. Fine-tuning is performed using 5-fold cross-validation on the OOD training set, starting either from the ArthroNat + flatbug model (blue) or from a control model trained on the COCO dataset (orange). Performance is evaluated on the OOD test set. Connecting lines indicate the mean value of each group, shown next to each box.

**Figure 8.**
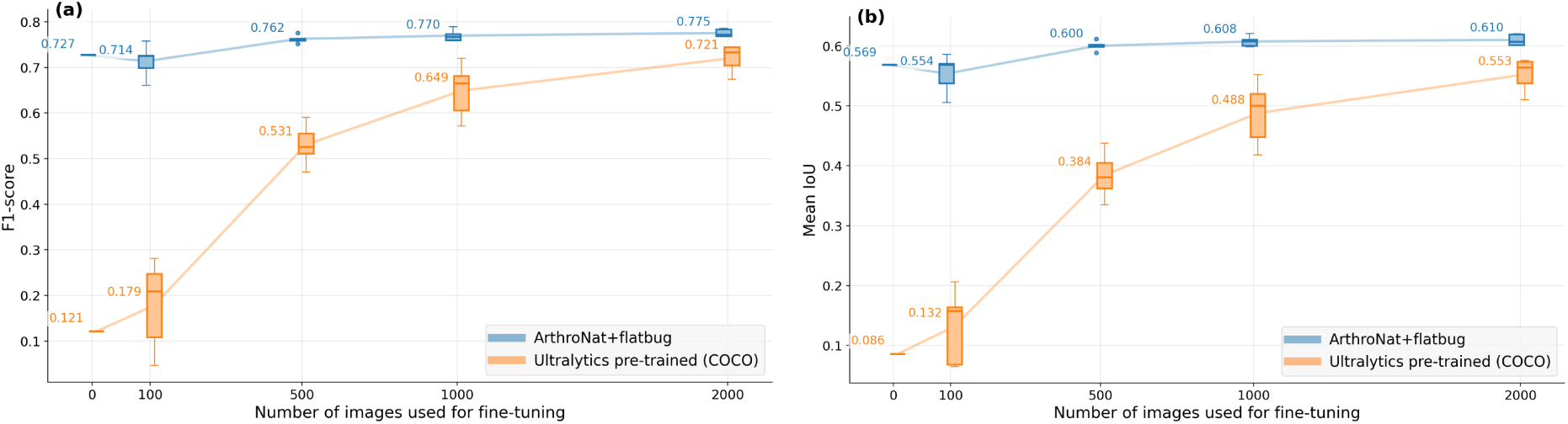
F1-score (a) and mean IoU (b) as a function of the number of images used for fine-tuning. Fine-tuning is performed using 5-fold cross-validation on the Lepinoc training set, starting either from the ArthroNat + flatbug model (blue) or from a control model trained on the COCO dataset (orange). Performance is evaluated on the Lepinoc test set. Connecting lines pass through the mean value of each group, shown next to each box.

## DISCUSSION

Arthropods are declining worldwide, making scalable biodiversity monitoring increasingly urgent. Automated arthropod detection is a key component of such monitoring, but remains constrained by the high cost of training-data annotation and the need for robust performance across diverse contexts and high taxonomic diversity. In that context, our main contributions are threefold: (i) we constitute a comprehensive detection dataset, encompassing almost 750 arthropod families, through an iterative annotation workflow, (ii) we release high-performing and deployable detection models, demonstrating strong detection performance as well as generalization capabilities, to new taxa and new application contexts, (iii) we propose a progressive set of adaptation strategies to transfer performance to new applications with increasing computational and/or data annotation investment.

Our first contribution is methodological and concerns dataset construction. Through our data-collection criteria, ArthroNat covers 749 terrestrial arthropod families found in Metropolitan France, providing an unprecedented level of family-level taxonomic diversity (Table 1). The iterative annotation process proved effective in constituting the dataset, with cross-validation showing that adding new batches consistently improved detection performance regardless of batch order^1^. This confirms the value of the iterative approach: as the model improves, its pre-annotations for subsequent batches become more reliable, making manual correction faster and less labor-intensive. This trend is consistent with similar iterative workflows reported in the literature (Tenckhoff et al., 2024). To determine the number of images and iterative annotation steps needed to build a robust detector (with an F1-score > 0.9), we observed that while a few thousand images were sufficient to reach acceptable arthropod detection performance (F1-score > 0.8), achieving F1-scores above 0.9 required substantially more data, with clear diminishing returns after the first thousand images. Similarly, mean IoU exceeded 0.8 only after nearly 14,000 images, and additional data may be needed to approach 0.9. These results align with studies reporting strong performance at dataset scales similar to ArthroNat (Mazen, 2023; Ștefan et al., 2025) whereas smaller datasets often remained below 0.8 on related metrics such as mean average precision (Chudzik et al., 2020; Sun et al., 2018), even with more than 2,000 images available. Although high F1-scores (>0.9) have also been reported with fewer than 1,000 images (Bjerge et al., 2025), that study included nearly 6,000 bounding-box annotations, suggesting that annotation density (boxes per image and total number of boxes) is a key determinant of model performance, in addition to the total number of images.

The broad taxonomic coverage of ArthroNat, encompassing 11 classes, 67 orders, and 749 families, allows our detection model trained on it to reach solid performance across arthropod taxa. First, considering taxonomic groups present in the ArthroNat dataset: The mean IoU remains consistently high (0.85 or above) across taxonomic levels, indicating robust bounding box quality regardless of taxon, whether looking at the taxonomic class– or order levels. While the two most represented arthropod classes (Insecta and Arachnida, 89% of the ArthroNat test set) show high F1-scores (≥0.84), some rarer classes have lower mean F1-scores. These remaining groups are less likely to be encountered in typical field observations, and several of the lowest-scoring taxa, including Branchiopoda, Entognatha, and the order Isopoda, are often small or cryptic and more commonly documented under magnification (e.g., with microscopes). As a result, their lower class-wise scores are expected to have limited impact on real-life field observations. We found no significant linear association between the number of images per taxon and model performance, indicating that taxon image count alone is not a reliable predictor of model performance.

Second, our model demonstrates a strong capability to generalize to taxa never seen during training. Performance assessments across different generalization levels show that the model maintains substantial detection capabilities (F1-score ≥ 0.9) even for completely unseen arthropod groups, highlighting the added value of taxonomic diversity in training data. Interestingly, performance at the same species level is sometimes lower than at higher taxonomic levels, which may be due to the larger and more diverse test set at the species level. This observation suggests that both within-species and cross-taxa generalization are important for evaluating model robustness in real-world applications. Although we demonstrate good generalization from European (French) arthropod families to those in South America, further testing is needed to assess generalization to other biodiverse regions, such as Africa, Oceania, and Asia.

Relative to generalization across datasets, our results show that transfer performance depends primarily on two controllable factors: training-data composition and the match between augmentation and target image structure. Across all five independent test sets, the model trained on ArthroNat+flatbug achieves the best overall performance, indicating that combining complementary datasets is the most reliable route to robust generalization. At the same time, performance remains dataset-dependent: scores are highest on datasets that are closest to the training distribution (SPIPOLL), and lower on less similar sets such as OOD and Lepinoc. This confirms that strong average performance does not remove the need for application-specific adaptation.

Beyond dataset composition, mosaic augmentation strongly influences transfer outcomes. When additional training data are not available, stronger mosaicing can recover much of the performance gap: on ArthroNat, OOD, and SPIPOLL test sets, an ArthroNat-only model trained with 3×3 mosaics performs close to the ArthroNat+flatbug model. Overall, mosaic augmentation beyond the default 2×2, especially 3×3 or 4×4 depending on the target dataset, can substantially reduce the performance gap caused by not including new data in training, such as flatbug. This aligns with studies demonstrating the effectiveness of mosaicing in improving small object detection and model robustness (Aldubaikhi & Patel, 2025; Dadboud et al., 2021). However, the optimal setting is not universal. On flatbug and Lepinoc, 4×4 mosaicing matches or exceeds 3×3, consistent with their smaller and denser bounding boxes. Together, these results indicate that adaptation should be guided by target image statistics, especially object apparent size and object density, rather than by a single default augmentation recipe. In parallel, object scale remains a key determinant of detection quality. On the ArthroNat test set, both F1-score and IoU increase with arthropod apparent size across all training scenarios, consistent with the broader object-detection literature (Liu et al., 2019). Differences between strategies are most visible at the size extremes: ArthroNat+flatbug and ArthroNat 3×3 remain strongest overall, while ArthroNat 4×4 gives the best IoU for the smallest decile but performs worse at the largest. Practically, performance curves plateau around an apparent size of about 10% of image area, suggesting a useful deployment target: whenever possible, acquisition protocols should aim for arthropods to occupy at least 0.1 of the image, as performance drops markedly below that threshold.

Based on these findings, we propose a progressive adaptation framework for new applications, ordered by increasing implementation cost. First, when the target context is reasonably similar to the training domain such as for the SPIPOLL application, the ArthroNat+flatbug model can be used directly and is the recommended baseline, as it provides the most consistent cross-dataset performance. Second, if initial performance is insufficient, augmentation should be tuned to target image properties, when training a new model. In particular, mosaic strength can be adjusted according to expected object size and crowding: 3×3 is a strong general option, while 4×4 may be preferable for very small or dense objects. Other relevant augmentations could indeed be chosen depending on the application (e.g. cutting and pasting arthropods out of natural images into uniform background if the application presents similar properties (H. Wang et al., 2021; Yun et al., 2021)). Third, if augmentation alone does not reach the required accuracy, adding external training data that are visually and structurally closer to the target application is the next lever, as illustrated by the gains from combining ArthroNat and flatbug. Finally, if the performance gap persists, targeted fine-tuning on target-domain annotations should be performed. In our experiments, fine-tuning with about 1,000 to 2,000 labeled instances improves both F1-score and mean IoU by roughly 5 percentage points. When annotation is expensive, this sequence minimizes manual effort; in a case where annotation would be relatively cheap, earlier fine-tuning may be preferable as it requires less training computation. Even in scenarios requiring such downstream fine-tuning, starting from our model remains more compelling than standard off-the-shelf pre-trained models, such as those pre-trained on the COCO dataset.

Our model and approach still face limitations, particularly in achieving F1-scores above 0.9 on generalization datasets like OOD and Lepinoc. In the specific case of Lepinoc, we identified that some shortcomings in the tiled data itself may have contributed to the lower performance metrics reported in the Supplementary Results. As the number of arthropod detection datasets continues to grow, we believe that combining a diverse range of datasets for training could further reduce this performance gap, an approach that has already shown promise in other studies (Svenning et al., 2025). Detection performance remains sensitive to the apparent size of arthropods in images, with reduced accuracy for small or distant individuals. While data augmentation techniques such as mosaicing can partially mitigate these effects, further research is needed to develop methods that improve robustness under these challenging conditions. We also see multiple parallels between our approach for constituting a diverse dataset and robust detector and approaches in arthropod image segmentation. Other pre-processing methods could further be explored, such as multi-scale tiling during both training and inference (Svenning et al., 2025), to reduce performance degradation in images with high arthropod density. Expanding datasets to include a broader range of imaging contexts, refining augmentation strategies, and exploring new model architectures may help address these limitations and advance the development towards universal, automated arthropod monitoring systems for ecological research and biodiversity assessment. This future work could also leverage our resources to improve detection in underrepresented contexts, expand to other regions or taxa, and expand from detection to robust object tracking for arthropods (Mirzaei et al., 2023; Pal et al., 2021).

## CONCLUSION

We present ArthroNat, a large-scale and taxonomically diverse dataset for French terrestrial arthropod detection, containing 13,661 images and covering no less than 749 families across the arthropod phylum. Trained on our dataset, our best arthropod detector achieves state-of-the-art performance (F1-score 0.91, mean IoU 0.83) on challenging natural background images. Our model generalizes well to previously unseen taxa and to diverse application settings. We also propose a stepwise adaptation framework for new domains, where performance can be improved using specific augmentations, adding complementary training data, and, when necessary, applying targeted fine-tuning. Our dataset, training and analysis code are publicly available on GitHub (github.com/edgaremy/arthropod-detection-dataset), and model weights are available on Hugging Face (huggingface.co/edgaremy/arthropod-detector), including both YOLO11l and lightweight YOLO11n variants for our two best training strategies.

With the expanding deployment of autonomous systems for non-destructive arthropod monitoring, the need for reliable data-processing tools is growing. We hope this work will contribute to making these tools widely accessible. Our model can help accelerate annotation of new detection datasets through model-assisted labeling, as achieved in this study. The lightweight model variants can be integrated into real-world monitoring systems, running on lightweight computers or embedded systems (Bjerge et al., 2021; Sittinger et al., 2024), creating a virtuous cycle of improved data processing and model performance.

Based on our conclusive results concerning generalization to unknown taxa, we would prioritize adding more annotated images of the same species, but shot using other methods, making the detectors more robust to new contexts. Such efforts will support large-scale biodiversity assessment and conservation, enabling automated monitoring in both citizen science and embedded camera trap applications (Serra-Marin et al., 2025), and ultimately advancing our ability to track and understand arthropod populations at unprecedented scales.

## ACKNOWLEDGMENTS

The authors would like to give special thanks to Lea Geng, Lowen Lecourt, Mathilde Llado and Melanie Jeanjean for their precious help in the manual annotation process, as well as Alice Dissoubray for her contributions with annotation as well as the validation process. We thank the NOÉ association (LEPINOC team) and the SPIPOLL program for providing the generalization datasets. In particular, we thank Jean Cohen for extracting a dataset with the highest possible diversity from the SPIPOLL database. We also thank the iNaturalist community for their contributions to the platform, which made this work possible. The authors also acknowledge the use of AI tools for facilitating code writing for handling data,plotting figures and for English language editing.

## SUPPLEMENTARY INFORMATION

### Additional details about the Data Collection

When we started this work, few publicly available datasets provided bounding boxes for a wide variety of arthropods. While knowing the exact species in an image is not required for detection, collecting data with species-level identifications can ensure the presence of arthropods in the images. Additionally, this approach allowed us to analyze performance based on taxonomic groups when interpreting our results. For these reasons, we decided to collect images with identified arthropod species on the iNaturalist platform. We only considered images with Research Grade Identifications. However, iNaturalist does not provide any bounding boxes for a detection task. This section describes how we selected the images to collect for our dataset, while the next section will elaborate on how we annotated the missing detection boxes.

**Table S1.**
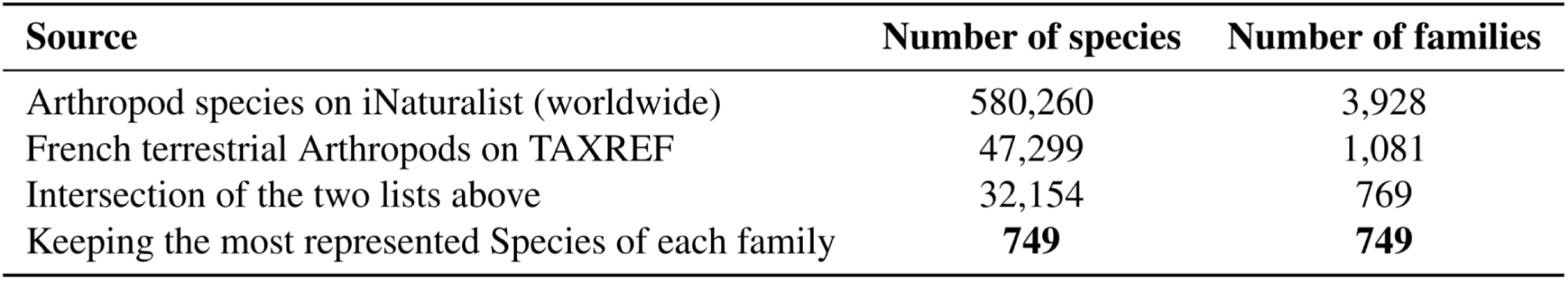
Summary of the scope-narrowing process for selecting the Arthropod species to include in our dataset.

| Source | Number of species | Number of families |
| --- | --- | --- |
| Arthropod species on iNaturalist (worldwide) | 580,260 | 3,928 |
| French terrestrial Arthropods on TAXREF | 47,299 | 1,081 |
| Intersection of the two lists above | 32,154 | 769 |
| Keeping the most represented Species of each family | **749** | **749** |

In the initial phase of dataset construction, we compiled a comprehensive list of arthropod species available on iNaturalist (as of January 2024), resulting in a dataset that would potentially include 580,260 species (worldwide, with research-grade data available). Though this amount of data certainly provided a wide array of taxonomic diversity, manually annotating such a vast quantity with reasonable resources was beyond reach. To address this challenge, we focused on a specific geographical region to reduce the number of species to a manageable size. We compiled a second list of arthropod species, containing only those found in Metropolitan France. This listing, indexed by TAXREF (Gargominy et al., 2020) (the official taxonomic reference for France), includes 47,299 species (as of January 2024) and excludes marine arthropods to focus solely on terrestrial and freshwater species from the continental region.

As iNaturalist was our only source of images for this dataset, we then intersected the two listings—iNaturalist’s global arthropod listing and the TAXREF listing for Metropolitan France—narrowing down the scope to 32,154 species. This intersection ensured that only arthropod species present in Metropolitan France and listed in iNaturalist are considered for further steps.

One goal was to ensure the data is as representative as possible of the wide diversity present among arthropods. To achieve this, we ensured that the dataset includes a similar number of images for each arthropod family. For each of these families, we identified the species with the most observations on iNaturalist. Note: It is important to distinguish between images and observations on iNaturalist. Each observation may include multiple pictures of the same species taken during a single event. This implies that collecting images without taking into account their observation id would result in weak data diversity: multiple images from the same observation are highly correlated and may not contribute to dataset diversity. We avoided this issue by using only one image per observation. We obtain a representative list spanning 769 distinct arthropod families, with each family represented by its most documented species. However, 20 of those families had no observations on iNaturalist, meaning we only have 749 arthropod families in reality. This narrowing allowed us to reduce the dataset to a manageable size for the subsequent annotation process, while still maximizing the diversity of the images collected. The process is summarized in Table S1.

Figure S1 and Figure S2 contain histograms of number of images per class, order and family in the final dataset as well as average, standard deviation, minimum and maximum number of images. Because of the scraping process, the number of images per family is mostly uniform, except for the few arthropod families with their most documented species having less than 20 distinct observations on iNaturalist. However, arthropod orders do not contain a uniform number of families, and classes do not contain a uniform number of orders. Thus the mostly uniform distribution of images at the family scale does not translate to a uniform distribution of images at the class and order levels.

**Figure S1.**
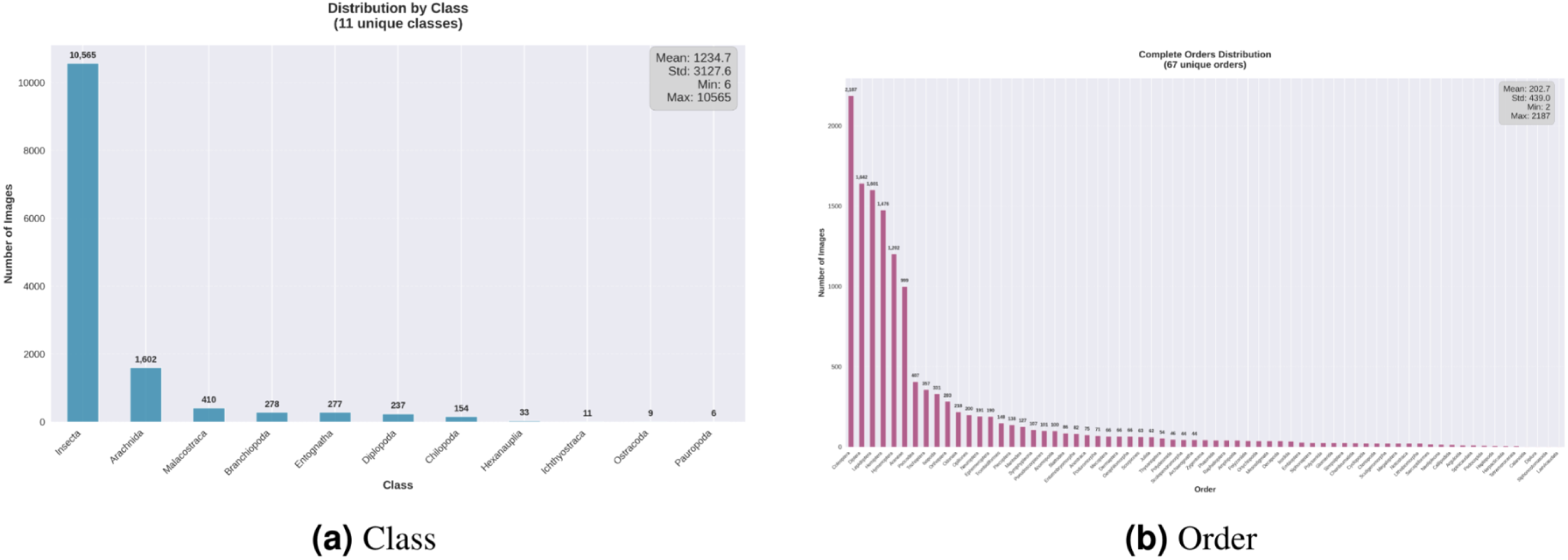
Histograms of the number of images per arthropod class (a) and order (b) in ArthoNat.

**Figure S2.**
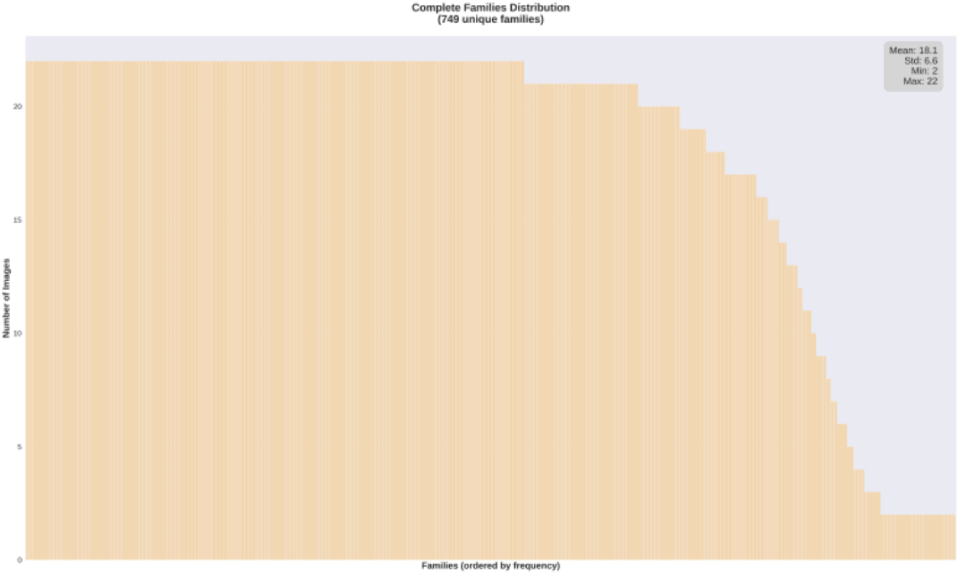
Histogram of the number of images per arthropod family in ArthoNat.

We display a few image samples of the ArthroNat dataset, with their labeled bounding box (see Figure S3). Thanks to our building process, ArthroNat is one of the arthropod detection datasets with the most diversity (see Table 1).

The final dataset is provided on GitHub (github.com/edgaremy/arthropod-detection-dataset), though due to copyrights, images are not directly stored there. Instead, we provide separately the labels and a script that download the images directly from their source link on iNaturalist.

**Figure S3.**
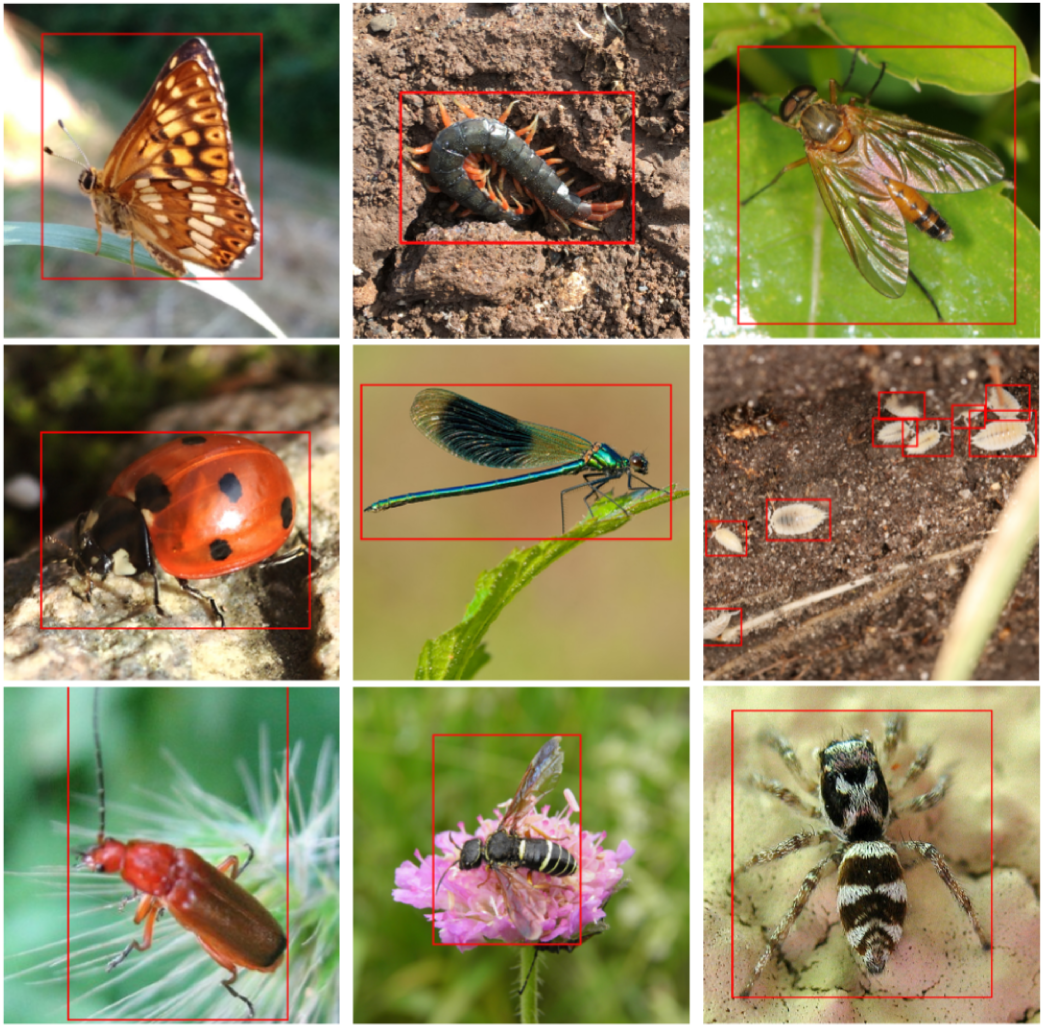
Visualization of ArthroNat images labeled with their detection bounding box. Arthropods have a strong diversity in shape and sizes, and can appear in varying numbers on images, with heterogeneous apparent size and diverse backgrounds.

### Details on the Iterative Annotation

**Figure S4.**
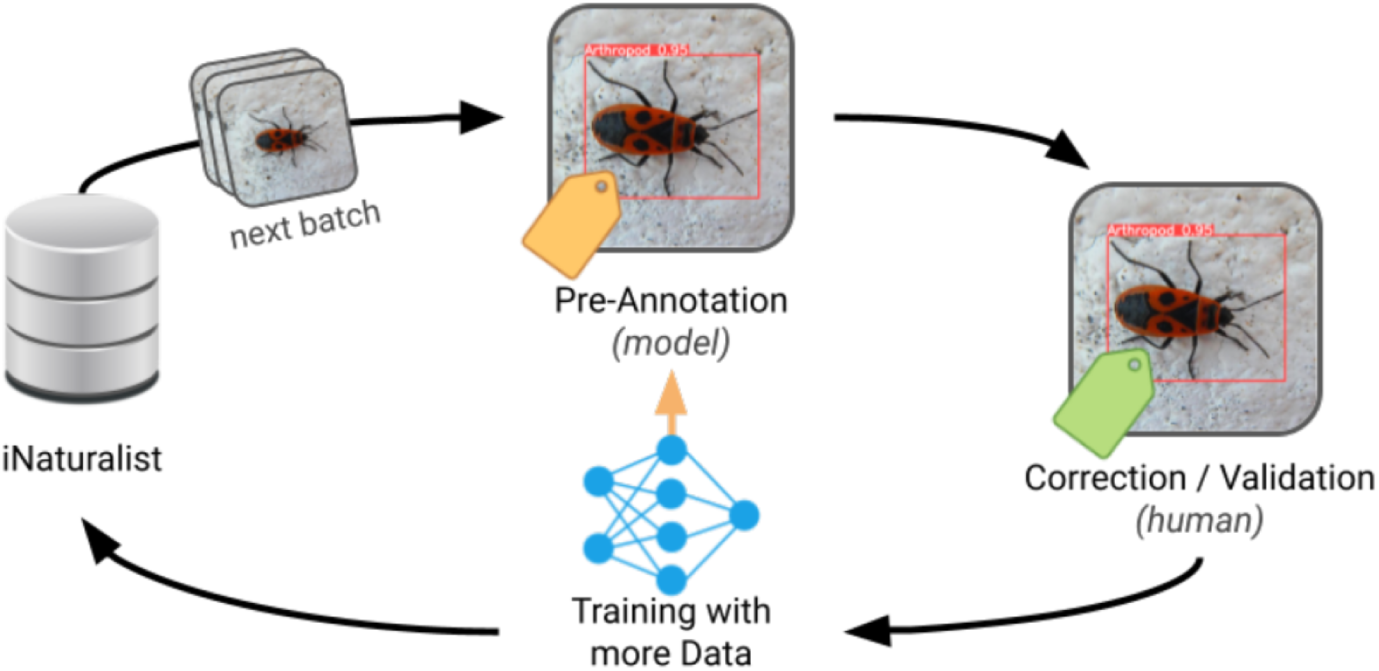
Iterative annotation process to obtain a detection dataset with proper bounding box labels.

For each iteration (see Figure S4), we fine-tuned a YOLO11 nano (YOLO11n) model using the Python library Ultralytics, starting from a baseline pre-trained on COCO (Lin et al., 2015). This baseline was further specialized by training on a custom wild bee segmentation dataset (Chiaburu et al., 2022), converted for object detection, instead of its original segmentation format. Although this baseline covers only wild bees, it provided a sufficiently relevant starting point for our insect detection task. For each subsequent image batch, this baseline model was fine-tuned on every previously annotated image. Keep in mind that this wild bee dataset was only used for the iterative process: final training done on our dataset starting from the baseline with COCO pre-trained weights.

### Per-order performance on the ArthroNat test set

**Table S2.** Order-level F1-scores and mean IoU of our baseline model on the ArthroNat test set (across all the 67 arthropod orders represented in Arthropod)

| Order | # images | F1-score | mean IoU |
| --- | --- | --- | --- |
| Coleoptera | 315 | 0.92 | 0.92 |
| Diptera | 251 | 0.86 | 0.92 |
| Lepidoptera | 220 | 0.94 | 0.91 |
| Hemiptera | 212 | 0.76 | 0.87 |
| Hymenoptera | 190 | 0.83 | 0.90 |
| Araneae | 144 | 0.91 | 0.92 |
| Psocodea | 64 | 0.78 | 0.86 |
| Trichoptera | 59 | 0.88 | 0.86 |
| Isopoda | 50 | 0.59 | 0.81 |
| Orthoptera | 38 | 0.87 | 0.84 |
| Ephemeroptera | 32 | 0.82 | 0.86 |
| Opiliones | 30 | 0.85 | 0.87 |
| Odonata | 29 | 0.98 | 0.90 |
| Neuroptera | 26 | 0.96 | 0.91 |
| Trombidiformes | 24 | 0.66 | 0.78 |
| Plecoptera | 23 | 0.81 | 0.93 |
| Pseudoscorpiones | 21 | 0.85 | 0.89 |
| Mantodea | 18 | 0.74 | 0.76 |
| Anomopoda | 17 | 0.83 | 0.88 |
| Symphyleona | 14 | 0.67 | 0.90 |
| Entomobryomorpha | 12 | 0.83 | 0.80 |
| Thysanoptera | 12 | 0.82 | 0.90 |
| Blattodea | 11 | 0.42 | 0.77 |
| Julida | 10 | 0.70 | 0.77 |
| Anostraca | 10 | 0.27 | 0.69 |
| Geophilomorpha | 9 | 0.78 | 0.79 |
| Scorpiones | 9 | 0.94 | 0.87 |
| Mecoptera | 9 | 0.89 | 0.93 |
| Dermaptera | 9 | 0.83 | 0.88 |
| Poduromorpha | 9 | 0.36 | 0.70 |
| Mesostigmata | 7 | 0.70 | 0.76 |
| Decapoda | 7 | 0.77 | 0.85 |
| Polydesmida | 7 | 0.93 | 0.83 |
| Amphipoda | 6 | 0.93 | 0.88 |
| Archaeognatha | 6 | 1.00 | 0.81 |
| Zygentoma | 6 | 0.59 | 0.73 |
| Scolopendromorpha | 6 | 0.91 | 0.84 |
| Sarcoptiformes | 6 | 0.45 | 0.92 |
| Raphidioptera | 6 | 1.00 | 0.89 |
| Onychopoda | 6 | 0.20 | 0.63 |
| Polyzoniida | 6 | 0.84 | 0.89 |
| Cyclopoida | 6 | 0.83 | 0.79 |
| Phasmida | 5 | 0.60 | 0.93 |
| Ixodida | 5 | 0.89 | 0.96 |
| Glomerida | 5 | 0.94 | 0.95 |
| Siphonaptera | 5 | 0.89 | 0.92 |
| Embioptera | 5 | 1.00 | 0.92 |
| Strepsiptera | 5 | 1.00 | 0.92 |
| Polyxenida | 4 | 0.89 | 0.91 |
| Ctenopoda | 4 | 0.48 | 0.89 |
| Chordeumatida | 4 | 1.00 | 0.86 |
| Lithobiomorpha | 3 | 1.00 | 0.96 |
| Tetramerocerata | 3 | 0.80 | 0.79 |
| Notostraca | 3 | 0.80 | 0.79 |
| Podocopida | 3 | 1.00 | 0.93 |
| Spinicaudata | 3 | 0.75 | 0.87 |
| Scutigeromorpha | 3 | 0.50 | 0.96 |
| Megaloptera | 3 | 1.00 | 0.95 |
| Harpacticoida | 2 | 1.00 | 0.94 |
| Arguloida | 2 | 1.00 | 0.93 |
| Callipodida | 2 | 1.00 | 0.95 |
| Neelipleona | 2 | 1.00 | 0.92 |
| Diplura | 1 | 1.00 | 0.82 |
| Laevicaudata | 1 | 1.00 | 0.63 |
| Haplopoda | 1 | 1.00 | 0.98 |
| Calanoida | 1 | 1.00 | 0.80 |
| Siphonostomatoida | 1 | 0.67 | 0.60 |
| Average on all 67 orders |  | 0.82 | 0.86 |

We computed the detection performance of our baseline detection model, averaged per order, on the ArthroNat test set (Table S2). The average performance of the 67 arthropod orders is also computed.

### Bounding box statistics of each main dataset

We computed bounding boxes statistics of each dataset used in this paper (Figure S5-S9), showing the distribution of bounding box count per image and bounding box apparent size. The mean value for both indexes can strongly vary from one dataset to the next.

**Figure S5.**
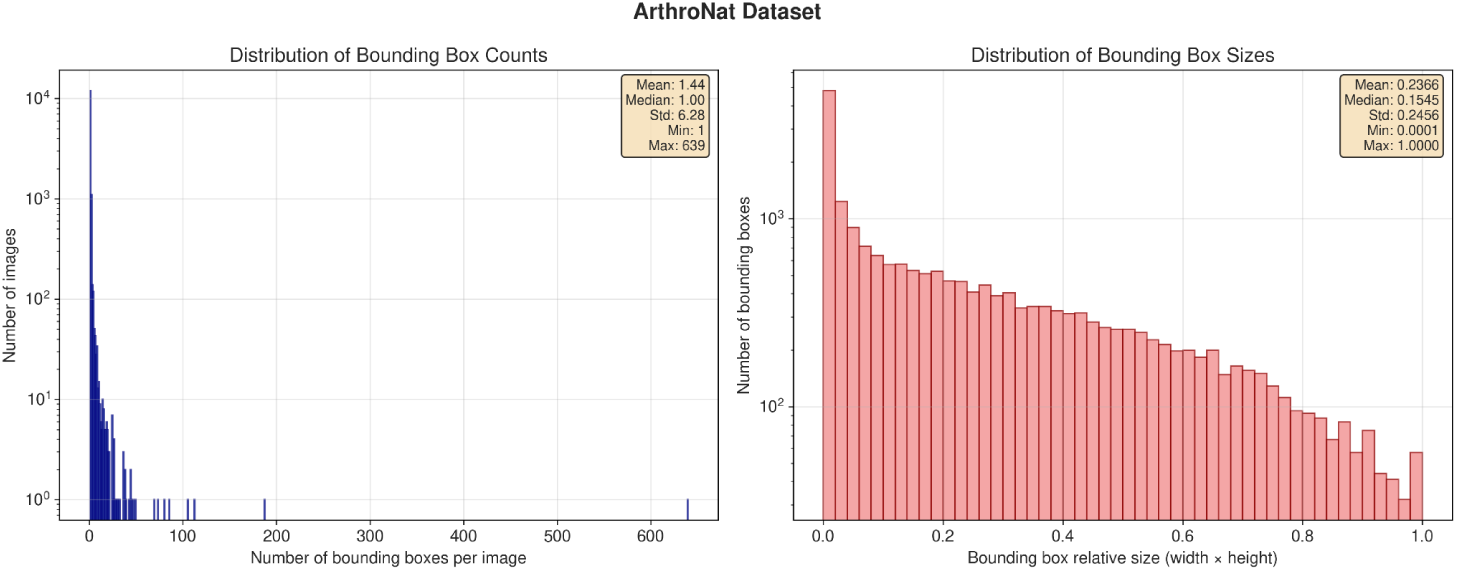
Distribution of bounding box count (left) and bounding box apparent size (right) within the ArthroNat dataset (all splits combined)

**Figure S6.**
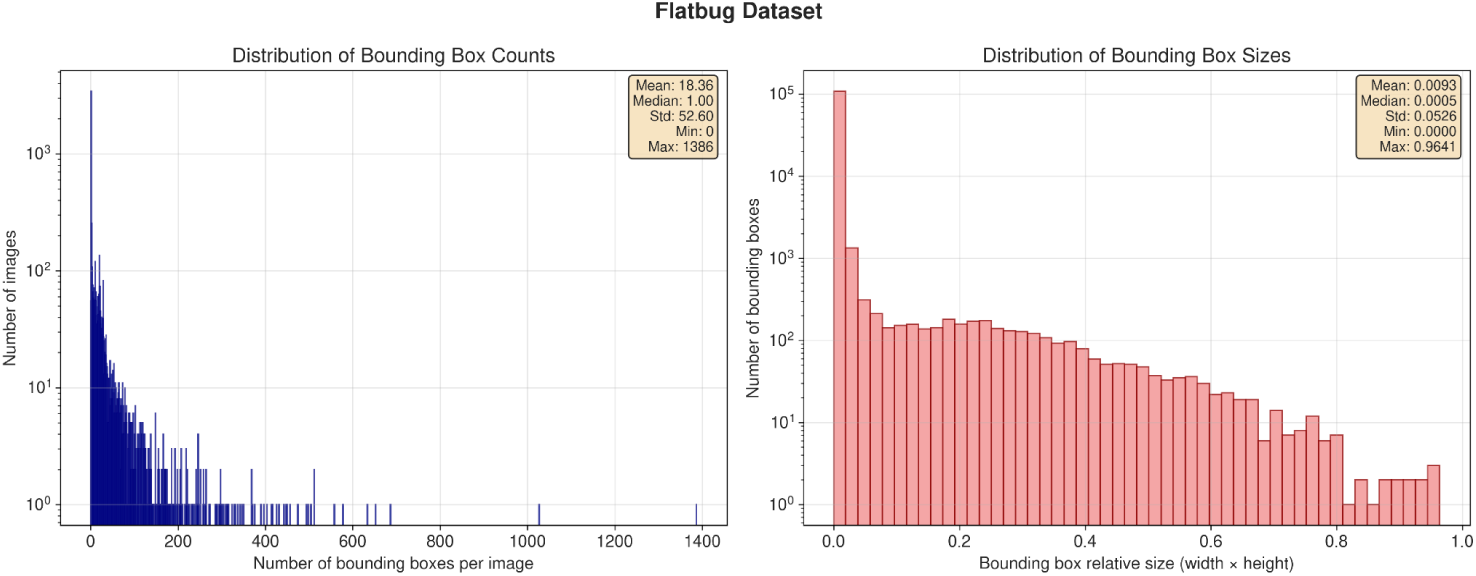
Distribution of bounding box count (left) and bounding box apparent size (right) within the flatbug dataset (all splits combined)

**Figure S7.**
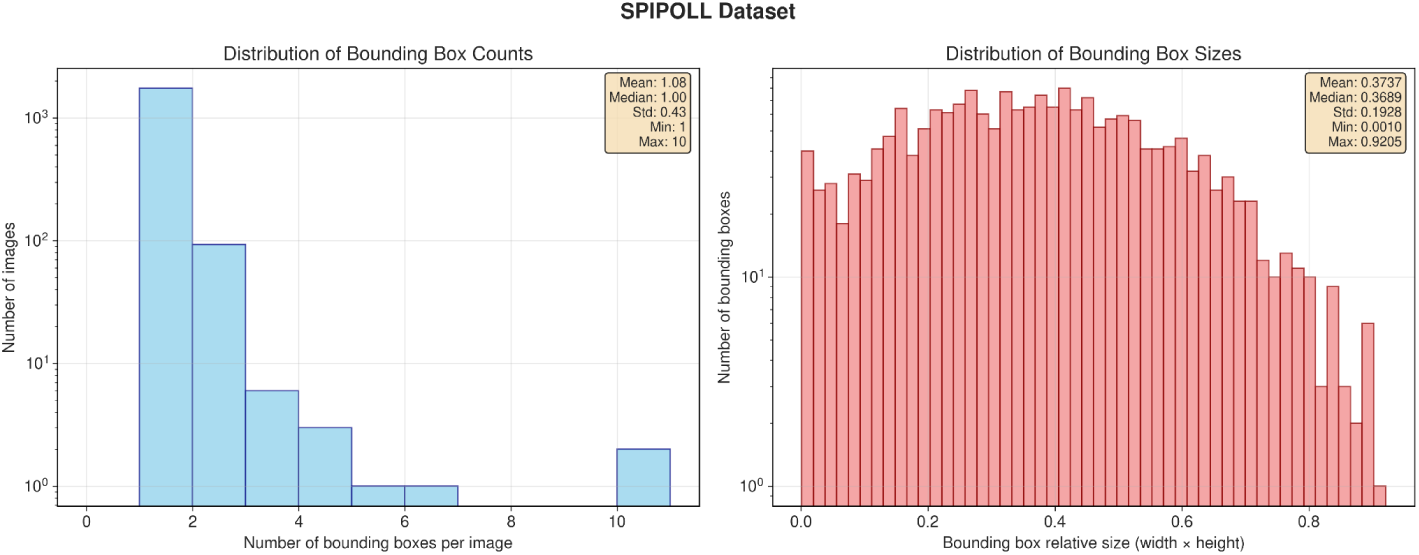
Distribution of bounding box count (left) and bounding box apparent size (right) within the SPIPOLL test dataset.

**Figure S8.**
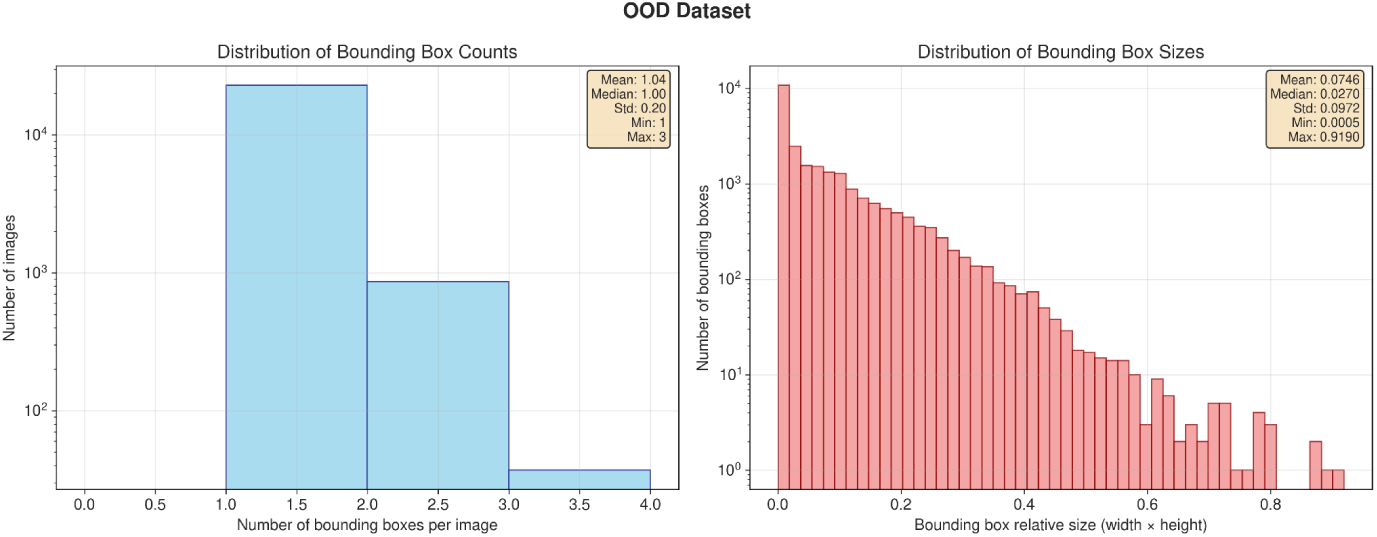
Distribution of bounding box count (left) and bounding box apparent size (right) within the OOD dataset (all splits combined)

**Figure S9.**
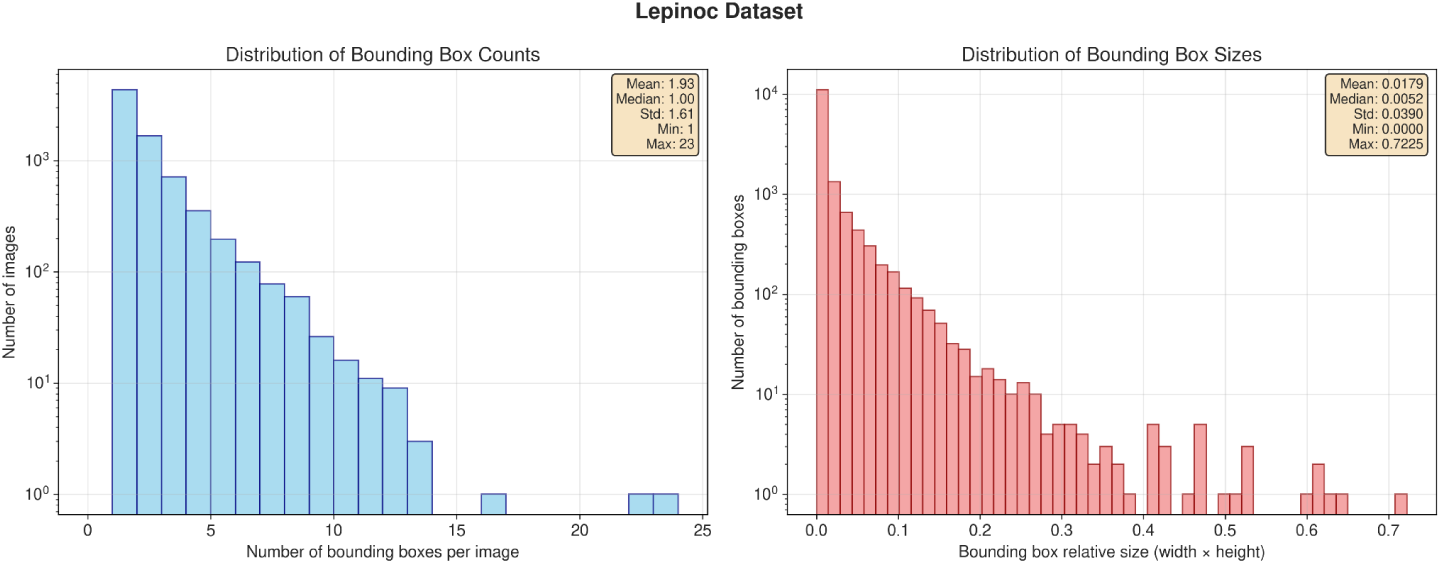
Distribution of bounding box count (left) and bounding box apparent size (right) within the LEPINOC dataset (all splits combined)

### Additional details on data used for across-dataset generalization

SPIPOLL is a French citizen science program designed to monitor plant–pollinator interactions, in which participants photograph insects visiting a given plant species during standardized 20-minute observation sessions. The images are then uploaded to an online platform, where they are collaboratively identified to the species or morphogroup level by the community. The program started in 2010 and in January 2025, it compiled 636,635 observations collected across 80,592 sessions, involving 548 insect taxa on 2,437 plant species and contributed by 4,947 participants. Our SPIPOLL test set was constructed to primarily capture variability among insect taxa, while also accounting for variation across plant species and participants. We retained all insect photographs that had received three independent validations. Each observation was assigned a weight combining two factors: the number of photographs associated with the same plant species and the number of photographs contributed by the same user. Greater weight was given to observations from less-sampled plant species, in order to prioritize plant diversity, while still accounting for user-level sampling effort. We then performed a weighted random sampling procedure: for each of the 548 insect taxa, a single observation was selected with a probability proportional to its weight. This procedure resulted in a final dataset comprising 548 photographs (one for each insect taxa), representing interactions with 473 plant species and contributed by 256 distinct users.

### Performance based on model size

We compared the performance of each YOLO11 model size, from *nano* up to *large*, on the ArthroNat test set (Figure S10). Each model is trained with the same configuration, still using the default parameter values provided by ultralytics’ library and notably the number of epochs set to 100. Though the smaller model (YOLO11n) already reaches above 0.8 on every metric used, each model size increase provides a small increase in performance.

**Figure S10.**
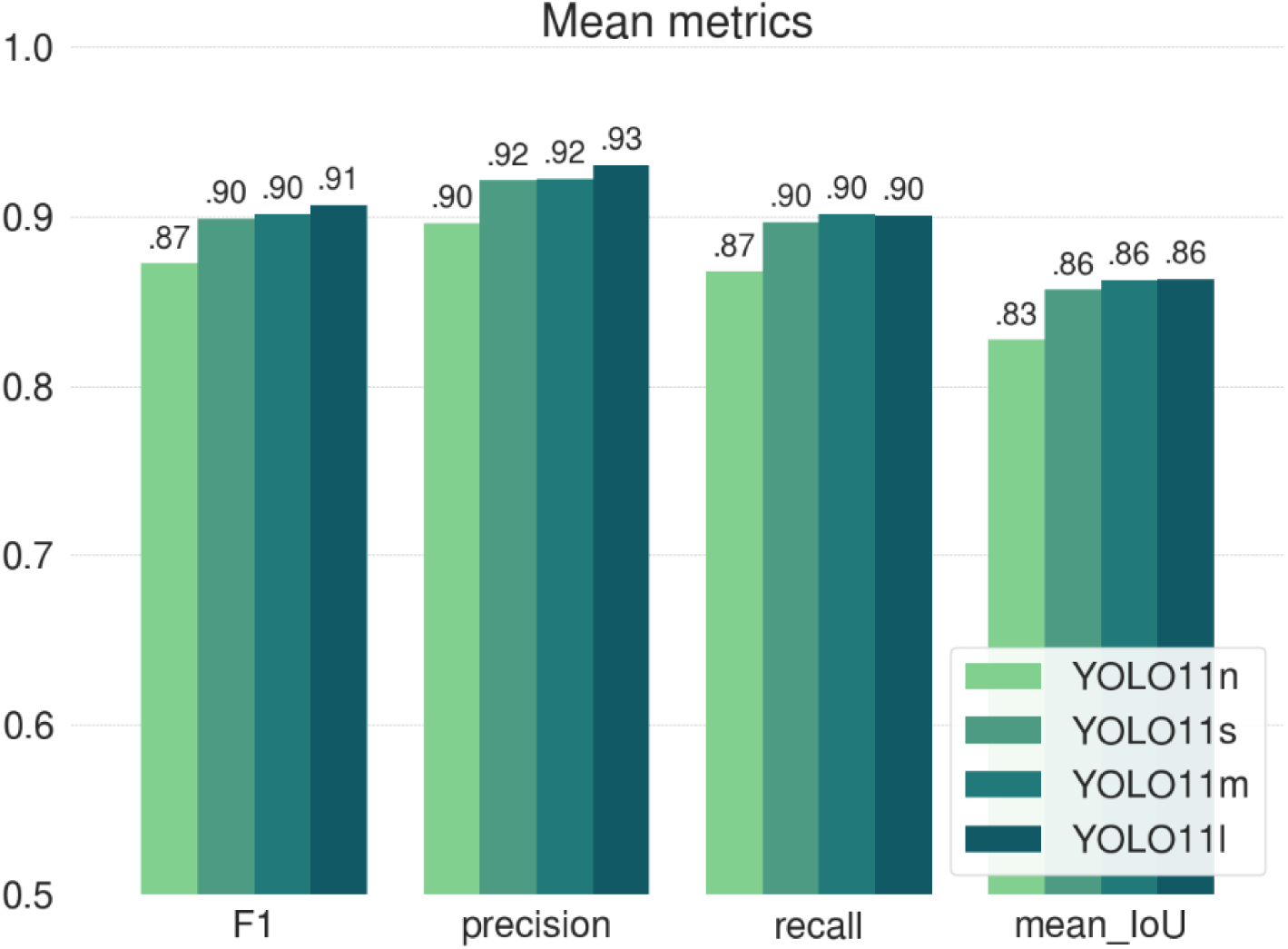
Mean metrics on the ArthoNat test set, depending on the detection model used. Each metric is computed by averaging the metric of each image.

To illustrate the computational cost against the performance gains, Figure S11 shows the average F1-score depending on the model size, in terms of number of parameters, or in terms of number of FLOPs. The models are trained and tested on ArthroNat. Both figures should give an idea of the potential loss of performance to be expected, going from a YOLO11l model to a smaller model of the family, which should also be similar for training scenarios including mosaicing augmentations, or training data from the flatbug dataset. Overall, the biggest leap in performance is observed between the nano (YOLO11n) and small (YOLO11s) variants. The improvements after that (from small to medium and from medium to large) are more incremental, with less return on performance invested. This suggests that the YOLO11s variant presents a good compromise between detection performance and computational cost, and that the nano variant should not be used, unless the computational requirements are very constraint (e.g. when deploying the model on a Raspberry Pi (Mathe et al., 2024)). The biggest variant we tested (large) remains the best choice if the objective is to maximize detection performance.

**Figure S11.**
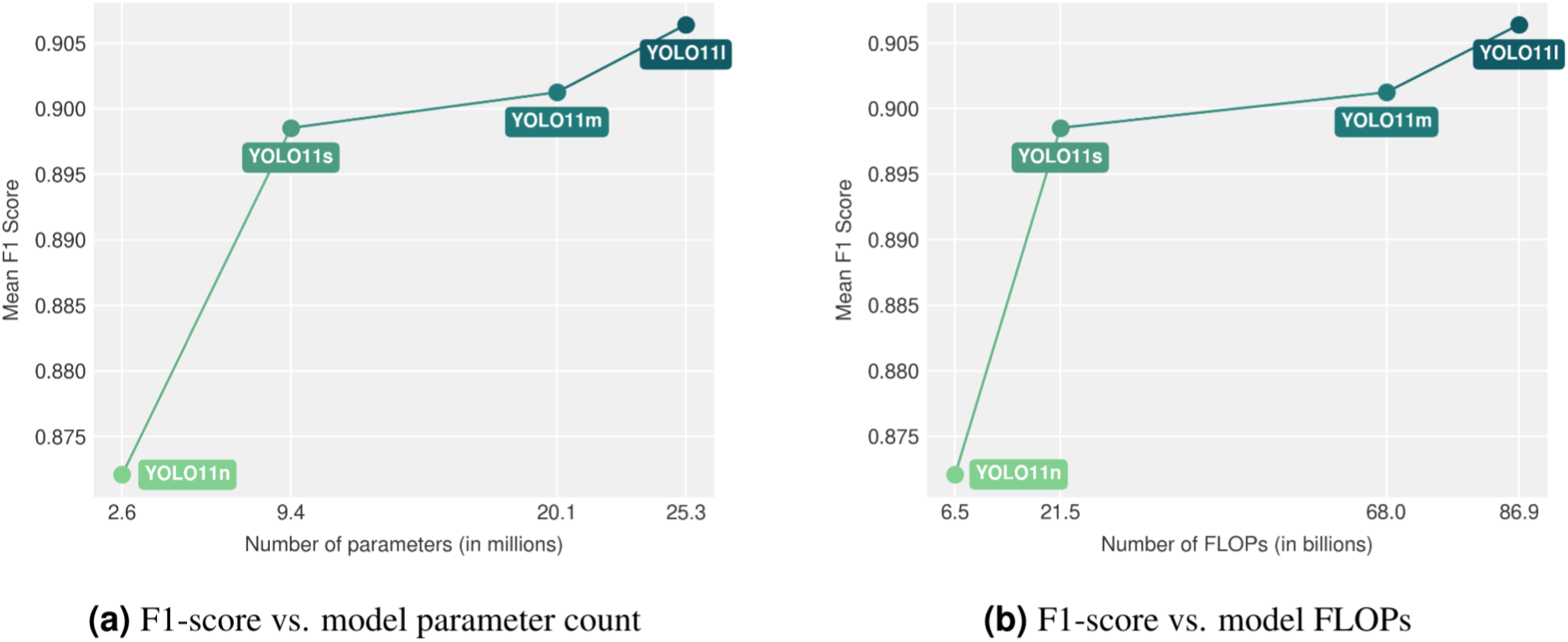
Mean F1-score on the ArthroNat test set, depending on the number of parameters of the model (a) and number of FLOPS (*floating-point operations per second*) (b)

### Mosaicing data augmentation

Mosaicing is a data augmentation technique that combines multiple images into a single training sample. This technique can add variability to the training data and help the model learn to detect objects in images with more diverse object densities and spatial arrangements.

**Figure S12.**
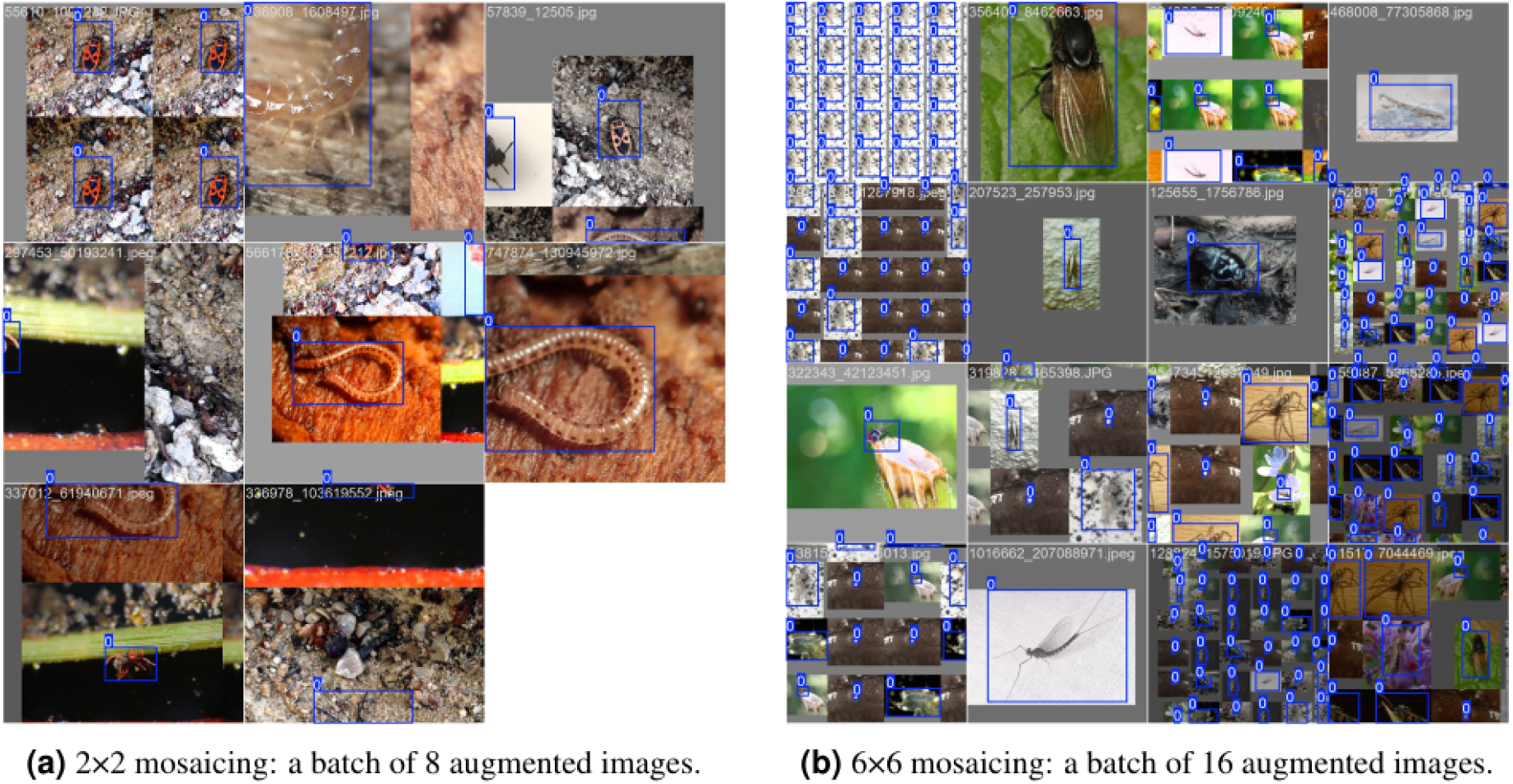
Examples of mosaicing data augmentation applied to arthropod detection. The technique combines multiple images containing arthropods into a single composite image, creating more challenging detection scenarios while preserving the bounding box annotations for each individual arthropod. Different grid sizes (from 2×2 to 6×6 in our implementations) provide varying scaling of arthropod apparent size relative to the image for training.

The mosaicing augmentation is applied with a configurable probability during training, allowing the model to experience both original images and augmented composite scenes. When an image is mosaiced, the size of the grid used is randomly chosen, to be smaller or equal to the maximum grid size defined for the training scenario. For instance, in Figure S12, we show examples of 2×2 and 6×6 mosaicing augmentations (potentially applied alongside other augmentations). Notice how the mosaic grid of each image in the batch varies respectively between 1×1 and 2×2 for the first example, and between 1×1 and 6×6 for the second example. Furthermore, the images chosen to create the mosaic can be multiple times the same image, or can be images picked randomly from a buffer of the entire dataset depending on a buffer parameter. In addition to the already provided 2×2 and 3×3 mosaicing, we implemented 4×4 and 6×6 augmentations, to push our testing further. Though we included 6×6 mosaicing in our experiments, the results were not conclusive and so were not included in the main text (with lower metrics than in any other training scenario depicted, this made 6×6 mosaicing a less favorable augmentation option to consider).

### On the shortcomings of the Lepinoc data and performance

We investigated why our various detection models would underperform on the Lepinoc dataset. As a reminder, the Lepinoc dataset consists of images with much higher resolution than the 640×640 input size of our YOLO models. To accommodate this, the original images were divided into a grid of 640×640 tiles, which form the basis of the dataset. However, the bounding box labels were manually annotated on the original full-size images, and the tile-level labels are derived by splitting each bounding box across the tiles it intersects. For cases where most of the bounding is on the same tile, this usually is no problem (see Figure S13). For many cases however, this implies the presence of divided bounding boxes, containing close to none of the arthropod itself (see Figure S14). This is due the the shape of arthropods, which can have long legs or antenna, making the bounding boxes contain mostly background: when the box is then divided across tiles, it should in many cases be reduced again in size, to properly fit the remaining parts of the arthropod. This adjustment was not applied when tiling the Lepinoc data, resulting in lower bounding box quality, and by extension, lower quality of performance assessment.

**Figure S13.**
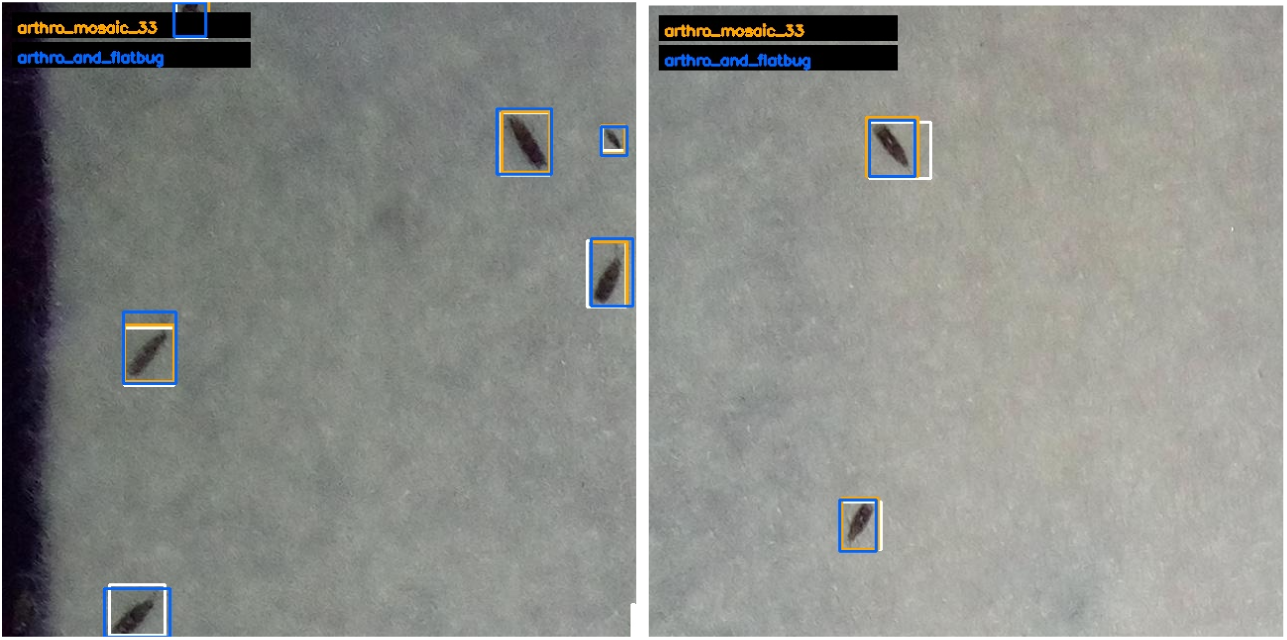
Examples of successful detections of our best models on the Lepinoc test set. White-stroke bounding box represents the ground-truth label.

**Figure S14.**
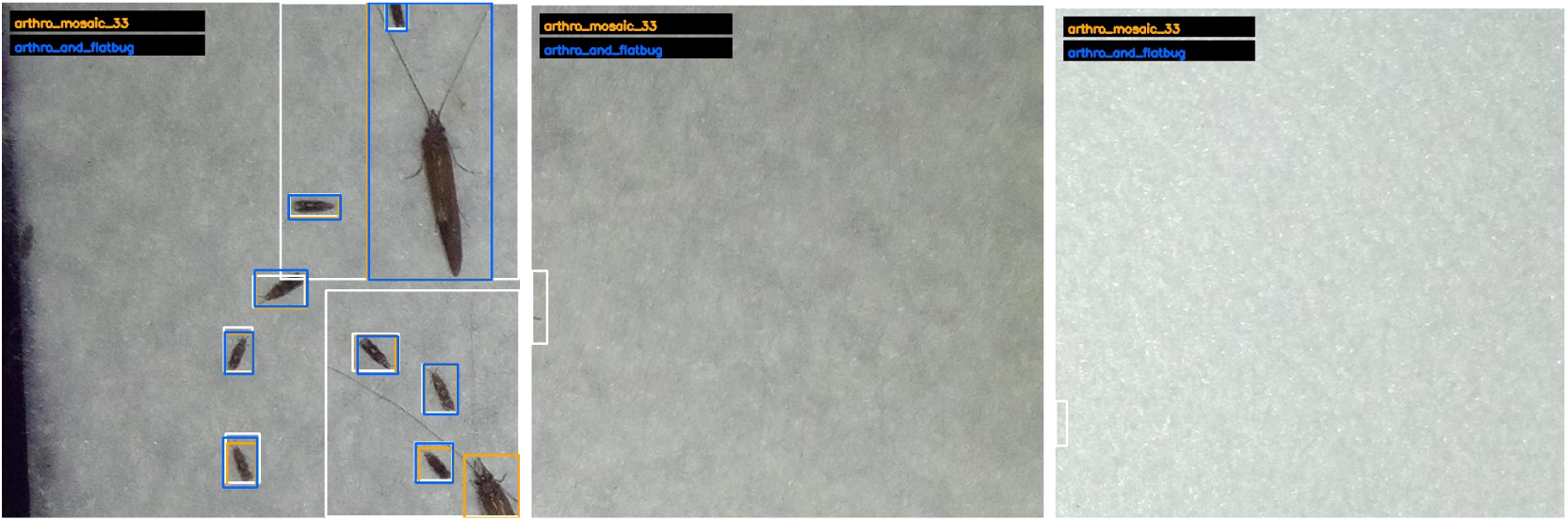
Example of tiled images in the Lepinoc dataset where certain arthropods are not properly contoured, or not easily visible, in both cases due to tiling. White-stroke bounding box represents the ground-truth label.

## Footnotes

1 The increase in variability of the results observable in Figure 4 can be explained by the progressive reduction of the test set size.

